# Involvement of ILC1-like innate lymphocytes in human autoimmunity, lessons from alopecia areata

**DOI:** 10.1101/2022.06.22.497177

**Authors:** Rimma Laufer Britva, Aviad Keren, Marta Bertolini, Yehuda Ullmann, Ralf Paus, Amos Gilhar

## Abstract

Here, we have explored the involvement of innate lymphoid cells-type 1 (ILC1) in the pathogenesis of alopecia areata (AA), because we found them to be significantly increased around lesional and non-lesional HFs of AA patients. To further explore these unexpected findings, we first co-cultured autologous circulating ILC1-like cells (ILC1lc) with healthy, but stressed, organ-cultured human scalp hair follicles (HFs). ILClc induced all hallmarks of AA *ex vivo*: they significantly promoted premature, apoptosis-driven HF regression (catagen), HF cytotoxicity/dystrophy and most important for AA pathogenesis, collapse of the HFs physiological immune privilege. NKG2D-blocking or IFNγ-neutralizing antibodies antagonized this. *In vivo*, intradermal injection of autologous activated, NKG2D+/IFNγ-secreting ILC1lc into healthy human scalp skin xenotransplanted onto SCID/beige mice sufficed to rapidly induce characteristic AA lesions. This provides the first evidence that ILC1lc suffice to induce AA in previously healthy human HFs *ex vivo* and *in vivo*, and further questions the conventional wisdom that AA is always an autoantigen-dependent, CD8+ T cell-driven autoimmune disease.

## Introduction

Alopecia areata (AA) is both the most common inflammatory hair loss disorder and one of the most common human autoimmune diseases and exerts a major negative impact on quality of life (Gilhar et al., 2012; Gilhar et al., 2019a; Korta et al., 2018; Pratt et al., 2017). Despite major recent advances in AA therapy, a causal therapy does not yet exist, and disease relapse after therapy discontinuation is the rule, not the exception in long-standing AA (Meah et al., 2020; Gilhar et al., 2019a). Thus, the currently available, purely symptomatic AA therapy, including JAK inhibitors (Gilhar et al., 2019b), remains unsatisfactory. Since the exact pathobiology of AA and its clinical variants remains to be fully characterized, the – likely diverse – disease-initiating factors that ultimately result in the characteristic AA hair loss pattern shared by all AA variants, require more comprehensive dissection for optimal, personalized therapeutic targeting (Bertolini et al., 2020; Paus et al., 2018).

Specifically, there is increasing awareness that a classical, autoantigen- and CD8+ T cell-dependent autoimmune variant of AA (AAA) and a possibly autoantigen-independent non-autoimmune variant (NAIAA) may have to be distinguished from each other (Gilhar et al., 2019a; Bertolini et al., 2020; Paus et al., 2018; Paus et al., 2020). This is in line with the long-standing, but often under-appreciated clinical recognition that AA shows a wide spectrum of phenotypes and sub-forms (Gilhar et al., 2012; Ikeda et al., 1965; Meah et al., 2021; King et al., 2022).

One reason why the currently available AA therapy is not entirely satisfactory may be related to as yet insufficient therapeutic targeting of innate immunocytes in the immunopathogenesis of human AA, namely in NAIAA, even though these are now recognized as major players in AA pathobiology (Ghraieb et al., 2018; Ito et al., 2008; Li et al., 2016; Uchida et al., 2020; Uchida et al., 2021).

Previously, we had demonstrated that AA lesions are associated with a massive increase in the number of perifollicular NKG2D+ NK cells (Gilhar et al., 2013a), which recognize the activating NKG2D ligand MICA, a “danger” signal that is greatly overexpressed by the epithelium of lesional AA hair follicles (HFs) (Ito et al., 2008; Li et al., 2016; Connell and Jabbari, 2022). Subsequent work has confirmed the key role of NKG2D and its activating ligands in human and murine AA (Xing et al., 2014; Petukhova et al., 2010). In fact, AA lesions can be induced experimentally in healthy human scalp skin *in vivo* by the transfer of interleukin 2 (IL-2)-activated NKG2D+cells (Gilhar et al., 2013a), most of which had NK cell characteristics, with only a small minority of CD8+ T-cells being present, i.e., the best-recognized pathogenic lymphocyte population in AA (Gilhar et al., 2012; Gilhar et el., 2013a; Pratt et al., 2017; Bertolini et al., 2020; de Jong et al.,2018). Moreover, pro-inflammatory mast cells (Bertolini et al., 2014) and (likely autoantigen-non-specific) γδ T-cells are also increased around/in lesional human AA HFs (Uchida et al., 2020). Finally, these “intermediate immunity” protagonists suffice to induce the hallmarks of AA *ex vivo* (Uchida et al., 2021).

Taken together, this questions whether pathogenic, autoreactive CD8+ T-cells are the only drivers of disease, and that all cases of AA, represent a genuine, autoantigen-dependent autoimmune disease (Bertolini et al., 2020; Paus et al., 2018) in the strictly defined sense of this term (Rose et al., 1993).

In our ongoing exploration of the role of innate/transitional immunity in the pathobiology of AA (Paus et al., 2020; Uchida et al., 2020; Uchida et al., 2021; Gilhar et al., 2019a; Bertolini et al., 2014), we therefore have asked in the current study whether innate lymphoid cells type 1 (ILC1 cells) (Zhou et al., 2020; Nabekura et al., 2021a; Colonna et al., 2018) can initiate human AA lesions.

We were interested in these immunocytes since human ILC1 cells secrete large amounts of interferon-γ (IFN-γ) (Ebbo et al., 2017), the crucial AA pathogenesis-promoting cytokine (Gilhar et al., 2012; Gilhar et al., 2019a; Paus et al., 2018), and this notably independent of classical autoantigen-specific CD8+ T-cell activities. These “unconventional” T-cells are placed in strategic tissue locations (Collins et al., 2017; Jiao et al., 2016; Kim et al., 2021) and represent an important link between innate and adaptive immunity (Vivier et al., 2018). While ILC1s play an essential role in human inflammatory bowel disease (IBD) (Ebbo et al., 2017; Luo et al., 2022; Clottu et al., 2022), their role in the pathophysiology of autoimmune hepatitis and rheumatoid arthritis requires further investigation (Ebbo et al., 2017; Fang et al., 2020; Yang et al., 2015), and their role in human autoimmune diseases overall remains insufficiently understood. We hypothesized that AA might offer a good model disease for interrogating this role.

ILC1 cells are classified as a component of type 1 immunity (Shannon et al., 2021), express NKG2D, recognize conserved phosphoantigens (Nabekura et al., 2021a), and contribute to immunity against tumor cells, e.g., through NKG2D activation (Dadi et al., 2016). The activating receptor NKG2D and its ligands (MICA, ULBP3) play an important role in innate (NK, ILC1), “translational” (γδ T-cells) and CD8 T-cell-mediated immune responses to tumors and in several autoimmune diseases (Frazao et al., 2019; Babic et al., 2018).

Given that ILC1 cells produce TH1-type cytokines (such as IFN-γ) and share several phenotypic markers with NK cells, namely NKG2D (Spits et al., 2016), it is challenging to distinguish NKs and ILC1 cells (Tulic et al., 2019; Zhang et al., 2018; Seillet et al., 2021; Conlon et al., 2021). The transcriptional and functional identity of ILC1 cells in humans is still a matter of debate, given that in contrast to other ILC subsets ILC1 cells seem to lack robust markers that enable their unequivocal identification and isolation (Bennstein et al., 2020). However, integrin α1 (CD49a) and integrin α2 (CD49b) have been used as two mutually exclusive markers to distinguish between NK and ILC1 cells, with NK cells being defined as CD49b+CD49a− and ILC1 as CD49b-CD49a+ (Gao et al., 2017; Vienne et al., 2021; Flommersfeld et al., 2021). Also, in contrast to ILC1 and ILC1-like cells, classical NK cells demonstrate high T-bet and Eomes expression (T-bet^hi^ /Eomes^hi^) (Verma et al., 2020). Therefore, for the purpose of this study, we define ILC1-like cells as CD49a+ CD49b- (Verma et al., 2020) and as lin-/CD127+/CD117-/CRTH2-phenotype, which are typical to classical ILC1 cells (Bennstein et al., 2020; Krabbendam et al., 2021), and also as T-bet^lo^/ Eomes^hi^ (Bennstein et al., 2020) (in contrast to classical T-bet^hi^/Eomes^lo^ ILC1 cells [Verma et al., 2020]).

Specifically, we have asked whether a) their number is increased in lesional AA skin, b) they can damage human HFs *ex vivo* in a manner that mimics the AA phenotype, and finally c) whether ILC1-like cells alone suffice to induce AA in previously healthy human scalp skin *in vivo*. To address these questions, we first analyzed the abundance, distribution and phenotype of ILC1-like cells in human AA skin lesions compared to healthy human control skin. We then co-cultured autologous ILC1-like cells with freshly organ-cultured scalp HFs from the same patient, i.e., under conditions where the epithelium of these HFs transiently undergo an acute stress response and overexpresses MICA (Uchida et al., 2021), to check whether these innate lymphocytes exert any HFs cytotoxicity and/or impact on the physiological immune privilege (IP) of HFs (Bertolini et al., 2020; Paus et al., 2005; Ito et al., 2004; Peters et al., 2007; Bertolini et al., 2016). Finally, we injected autologous ILC1-like cells intradermally into healthy human scalp skin xenotransplants from the same human volunteers on SCID/beige mice to probe whether this suffices to induce classical AA hair loss lesions *in vivo*.

Taken together, our data show that ILC1-like cells are increased in AA lesions and suffice to induce an AA phenotype in healthy human HFs *ex vivo* and *in vivo*. This provides the first functional evidence of a key role of ILC1-like innate lymphocytes in a model human autoimmune disease (Colonna et al., 2018; Seillet et al., 2021; Conlon et al., 2021; Flommersfeld et al., 2021; Daussy et al., 2014; Park et al., 2019) - but also questions whether AA always a classical autoimmune disease is and underscore the role of innate immune cells in AA pathobiology.

## Results

### Peri- and intrafollicular infiltrates of ILC1-like cells are seen in both lesional and non-lesional alopecia areata (AA) skin

First, we investigated whether healthy and AA-affected human skin differ in their content and/or distribution of ILC1-like cells, using a comprehensive set of triple-immunofluorescence (IF) staining best suited to identify these immunocytes (Seillet et al., 2021; Bennstein et al., 2020; Gao et al., 2017). This revealed the presence of only extremely few ILC1-like cells in healthy control skin with all three staining settings employed (Eomes+, CD49a+, NKG2D+ [**Figure 1A and Figure 1----figure supplement 1A**], Eomes+,c-KIT−, CD49a+ [**Figure 1B and Figure 1----figure supplement 1A**], or NKp44+, CD103+, T-bet-cells [**Figure 1C and D and Figure 1----figure supplement 1A**]) (Kim, 2015; Fuchs et al., 2013; Salimi and Ogg, 2014). These cells appeared to be preferentially scattered along the papillary dermis of healthy scalp skin biopsies and around the HFs (**Figure 1C**). This is reminiscent of the few Vδ1+T-cells detectable in healthy human skin that also have a preferential perifollicular location and may “police” the skin for molecular indications of tissue stress, namely of HFs (Uchida et al., 2020; Uchida et al., 2021).

**Figure. 1.**
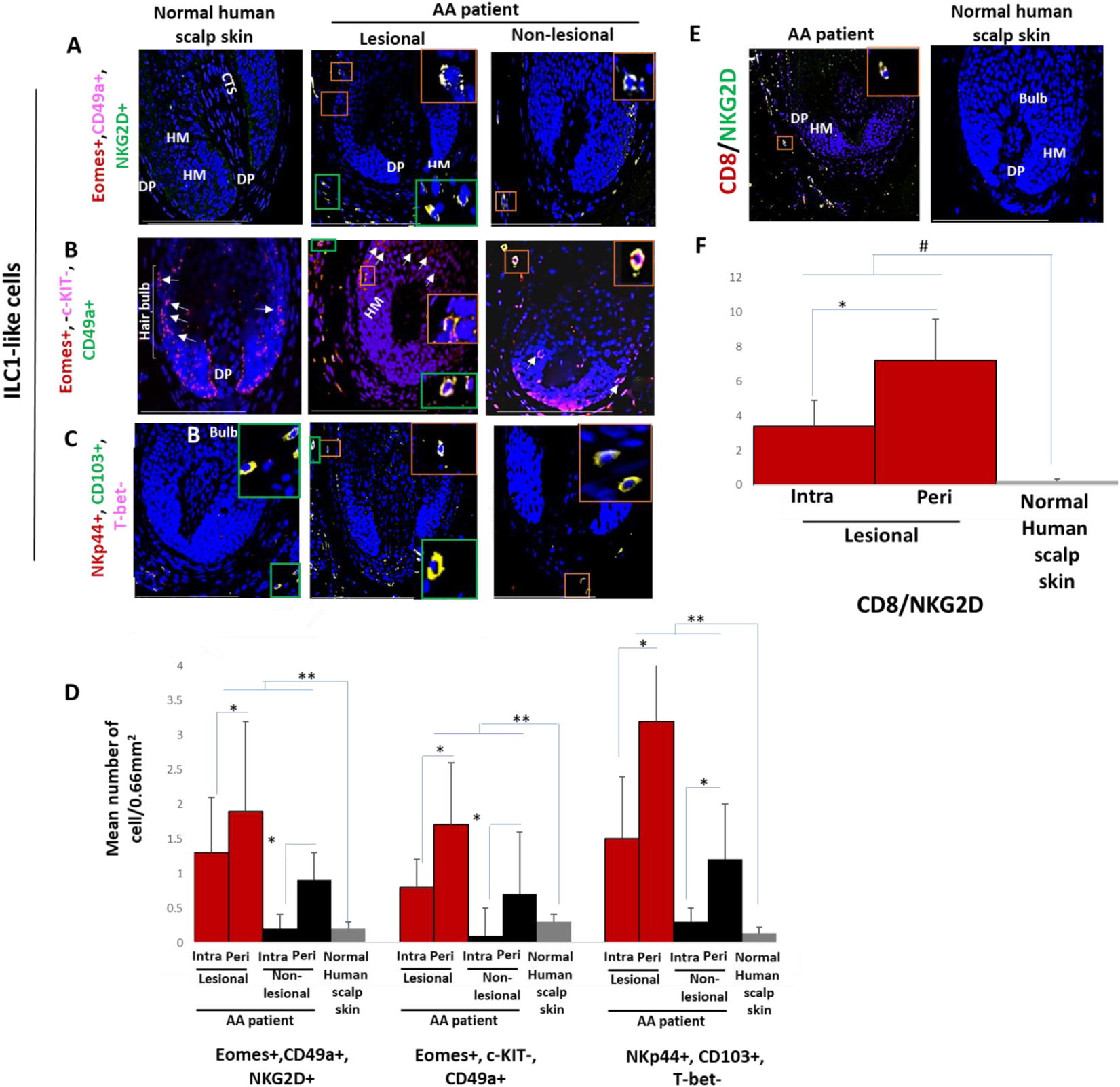
Immunofluorescence microscopy analyses of ILC1-like and CD8+/NKG2D+ cells in AA scalp skin. (**A**) ILC1-like cells (EOMES+, CD49a+ and NKG2D+) around HF in normal scalp skin, intrafollicular and perifollicular ILC1-like cells infiltrates in lesional and in non-lesional AA scalp patient.(**B**) EOMES+,c-KIT−,CD49a+ and (**C**) NKp44+, CD103+, T-bet-ILC1-like cells. (**D**) Quantitative immunohistomorphometry (qIHM) shows increased number of ILC1-like cells in AA patients as compared to normal volunteers and increased number of the cells in lesional versus non-lesional areas of the patients. There is a significant increased perifollicular than intrafollicular ILC1-like cells in the lesional and non lesional areas. (**E**) CD8+/NKG2D+ cells around HF in AA scalp patient and absence of these cells in normal scalp skin of normal scalp skin. (**F**) There is an increased number of CD8+/NKG2D+ cells in HFs of AA patients compared to normal scalp skin and a significant lower number of ILC1-like cells versus CD8+/NKG2D+ cells in AA scalp skin. N=6 biopsies /AA patients and 6 biopsies /healthy donors from 6 independent donors, 3 areas were evaluated per section, and 3 sections per biopsy. Following Shapiro-Wilk test, Student’s *t*-test: *\*p* < 0.05, \*\**p* < 0.01 or Mann Whitney *U* test: ^#^*p* < 0.05. Scale bars, 50 µm. **CTS** - connective tissue sheath, **DP** - dermal papilla, **HM** - hair matrix, White arrow-c-KIT stained melanocyte.

Instead, intra and peri-follicular infiltrates of ILC1-like cells were frequently present in lesional AA HFs (**Figure 1A, B, C and D and Figure 1----figure supplement 1A**), typically in conjunction with a dominant infiltrate of CD8+/NKG2D+ cells around the hair bulb (p<0.05) (**Figure 1E and F**). Importantly, the number of ILC1-like cells was already significantly increased in/around non-lesional AA HFs compared to healthy scalp skin (p<0.01) (**Figure 1A, B, C and D and Figure 1----figure supplement 1A**). This may indicate that ILC1-like cells may actually have arrived around the HFs before the CD8 cells and may have contributed to attracting the CD8 cells into the perifollicular space.

This strongly suggested that ILC1-like are not mere bystanders attracted only secondarily to the HFs by CD8 T-cells, similar to, but more pronounced than we have recently observed regarding perifollicular Vδ1+T-cells in non-lesional AA skin (Uchida et al., 2020). This invited the hypothesis that ILC1-like cells are actively involved in transforming healthy human scalp HFs into lesional AA HFs.

### T-bet^lo^/Eomes^hi^ ILC1-like cells can be expanded from human peripheral blood mononuclear cells (PBMCs) in vitro

To functionally probe this hypothesis, we isolated, purified and characterized human peripheral blood-derived ILC1-like cells as the most suitable cell source for the planned HF-immunocyte co-culture studies. The scarcity of ILC1-like cells in healthy human skin, compared to their relative abundance in peripheral blood (Colonna et al., 2018; Artis and Spits. 2015) necessitated to isolate autologous ILC1-like cells from the latter source rather than from skin (Teunissen et al., 2014). To facilitate ILC1-like cells isolation, PBMCs of healthy volunteers were first cultured with high-dose IL-2 (100 U/mL) in the presence of IL-18 (1 µg/1 ml) IL-33 (1.5 µg/5 ml) and IL-12 (1.5 µg/5 ml), since these cytokines induce ILC1-like cells expansion (Salimi and Ogg, 2014; Silver et al., 2016; Orimo et al., 2020; Ohne et al., 2016). When ILC1-like cells were sorted by FACS Aria and characterized by FACS analysis on day 7 of culture, low T-bet and high Eomes expression were observed (**Figure 2A**), in contrast to classical T-bet^hi^ and Eomes^lo^ ILC1 cells (Jiao et al., 2016; Vivier et al., 2018; Zhang et al., 2018). In addition, the ILC1-like cells expressed and shared the following markers with classical ILC1 cells: LIN-CD3/CD1a/D14/CD19/CD34/CD123/CD11c/BDCH2/FcεR1α/TCRαβ/TCRγδ/CD56), CD127+, CD161+, c-KIT− and CRTH2- (Zook and Kee, 2016; Bernink et al., 2017; Simoni and Newell, 2017) (**Figure 2A**). In contrast to NK cells, ILC1-like cells also expressed the expected high levels of integrin α1 (CD49a), combined with the absence of integrin α2 (CD49b) (Jiao et al., 2016) (**Figure 2A**). All these characteristic markers of ILC-like cells were absent in the control unstimulated PBMCs (**Figure 2B and C**).

**Figure. 2.**
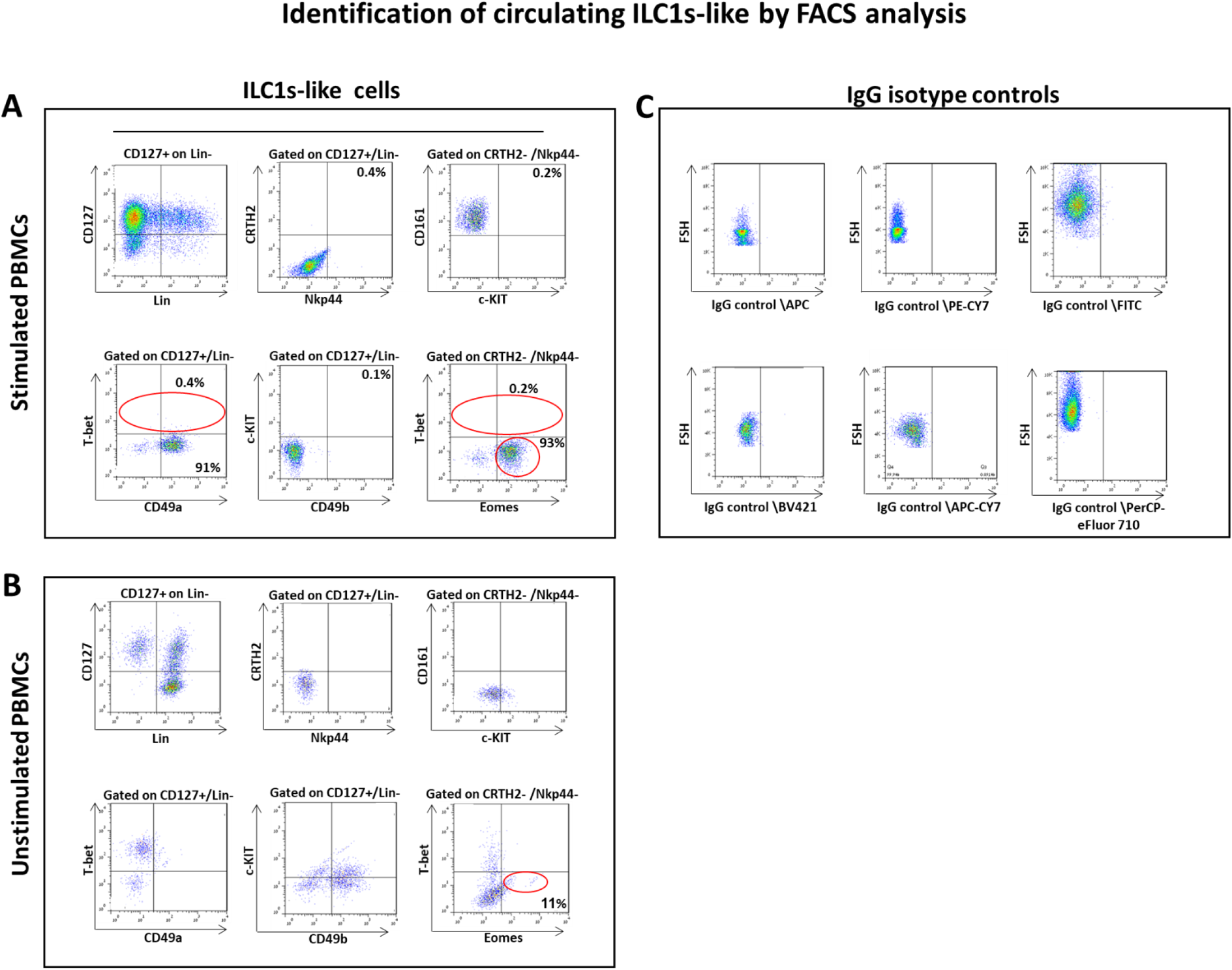
Circulating ILC1-like cells expanded and characterized by FACS analysis. (**A**)PBMCs activated by IL-18, IL-33 and IL-12 were sorted by FACS Aria and characterized by FACS analysis. ILC1-like cell markers were identified by the expression of CD127+, CD161+, c-KIT− and CRTH2-, high levels of integrin α1 (CD49a) expression, combined with the absence of integrin α2 (CD49b) and transcription factors Eomes^hi^ and T-bet^lo^ (**B**) unstimulated PBMCs (**C**) isotype controls. N=10 blood donors, 1.5×10^6^ cells/blood donor, analysis was performed in triplicates from each of the blood donor. Following Shapiro-Wilk test, Student’s t-test, *p*<0.05.

This immune phenotype suggests that the immune cells used in our study are best classified as ILC1-like cells (Nabekura et al., 2021a), and documents that all experiments reported below were indeed performed with autologous ILC1-like cells rather than with NK cell subpopulations. Note that we had previously shown that NKG2D+/CD56+ NK cells suffice to induce AA lesions in human skin *in vivo* (Gilhar et al., 2013a; Laufer Britva et al., 2020) while iNKT cells are AA-protective in the humanized AA mouse model (Ghraieb et al., 2018). Subsequently, these ILC1-like cells were either used for HF co-culture assays or injected into healthy human scalp skin xenotransplants on SCID/beige mice (Gilhar et al., 2013a; Ito et al., 2005a). As controls, we also isolated ILC2 and ILC3 cells, which failed to induce AA phenotype in a sharp contrast to the ILC1-like cells (see **Materials and Methods)**.

### ILC1-like cells induce HF cytotoxicity *ex vivo*

Next, we functionally probed the interaction of ILC1-like cells with HFs that were investigated here as a model human (mini-)organ in which the interactions of a healthy human tissue system with defined, autologous immunocyte populations can be interrogated *ex vivo* in the absence of any confounding systemic immune or neural inputs (Uchida et al., 2021). For this, microdissected, organ-cultured human scalp HFs (Langan et al., 2015) were co-cultured for six days with autologous, peripheral blood-derived, purified, IL-12/IL-18/IL-33-prestimulated ILC1-like cells, or with autologous human CD8+NKG2D+ cells (=positive control), ILC2, ILC3 cells, or PBMCs non-specifically activated with PHA (PBMCs/PHA) (=negative controls).

Importantly, only scalp HFs in the anagen VI stage of the hair cycle were used (identified as described) (Kloepper et al., 2010) that had been freshly placed into HF organ culture for 24 hours, since these HFs are maximally “stressed,” in contrast to non-cultured HFs, i.e. immediately after isolation, or that had already undergone several days of adjusting to the harsh conditions of serum-free organ culture (Uchida et al., 2021; Langan et al., 2015). These “stressed” day 1 HFs temporarily up-regulate MHC class Ia and ß2-microglobulin while the expression of IP guardians, i.e. αMSH and TGFβ2 remains unchanged (Uchida et al., 2021), indicating a transiently weakened, but partially maintained HF immune privilege (Bertolini et al., 2020; Ito et al., 2004). The expression of molecules associated with tissue stress, i.e. the intrafollicularly produced neurohormone, CRH (Ito et al., 2005b), and the NKG2D ligand MICA/B is also higher in day 1 organ-cultured HFs compared to freshly microdissected HFs or after day 3 of organ culture. Day 1 HFs also show signs of mild HF dystrophy (as evidenced by increased lactate dehydrogenase [LDH] release into the medium), and express chemokines recognized for their relevance in AA pathobiology, i.e. CXCL10 and CXCL12 (Uchida et al., 2021; Ito et al., 2020). Thus, day 1 HFs are ideally suited for interrogating human immunocyte interactions with a transiently “stressed”, but otherwise healthy human (mini-)organ that overexpresses the NKG2D-activating “danger” signal, MICA/B, under physiologically relevant *ex vivo* conditions (Uchida et al., 2021; Langan et al., 2015).

First, we studied the cytotoxic effects of ILC1-like cells on healthy human scalp HF *ex vivo* by measuring the HF release of LDH into the culture medium. This not only showed significantly higher LDH release induced by ILC1-like cells than by co-culture with all three negative control cell populations (ILC2s, ILC3s or PBMCs/PHA) but also even higher HF cytotoxicity levels than those induced by CD8+/NKG2D+ cells (p<0.01), namely after three days of co-culture (**Figure 3**). These HF cytotoxicity results were fully corroborated by characteristic morphological signs of HF dystrophy following co-culture with ILC1-like cells; while CD8+/NKG2D+ cells induced similar dystrophy phenomena, these were not seen after co-culture with PBMC/PHA (**Figure 4A, B and C**). The induction of significant HF dystrophy by ILC1-like cells *ex vivo* was further documented by the presence of pathological melanin clumping and ectopically located intrafollicular melanin granules (Bodó et al., 2007; Hendrix et al., 2005) (**Figure 4D, E, F and G**) and by decreased proliferation and increased apoptosis of hair matrix keratinocytes (**Figure 4H, I, J and K**). Both was also seen in the CD8+/NKG2D+ group (positive control), but not in HFs co-cultured with PBMCs/PHA (negative control) (p<0.001, p<0.01 respectively). Thus, autologous ILC1-like cells alone suffice to induce substantial HF cytotoxicity *ex vivo* if co-cultured with transiently “stressed”, but otherwise healthy human scalp HFs.

**Figure. 3.**
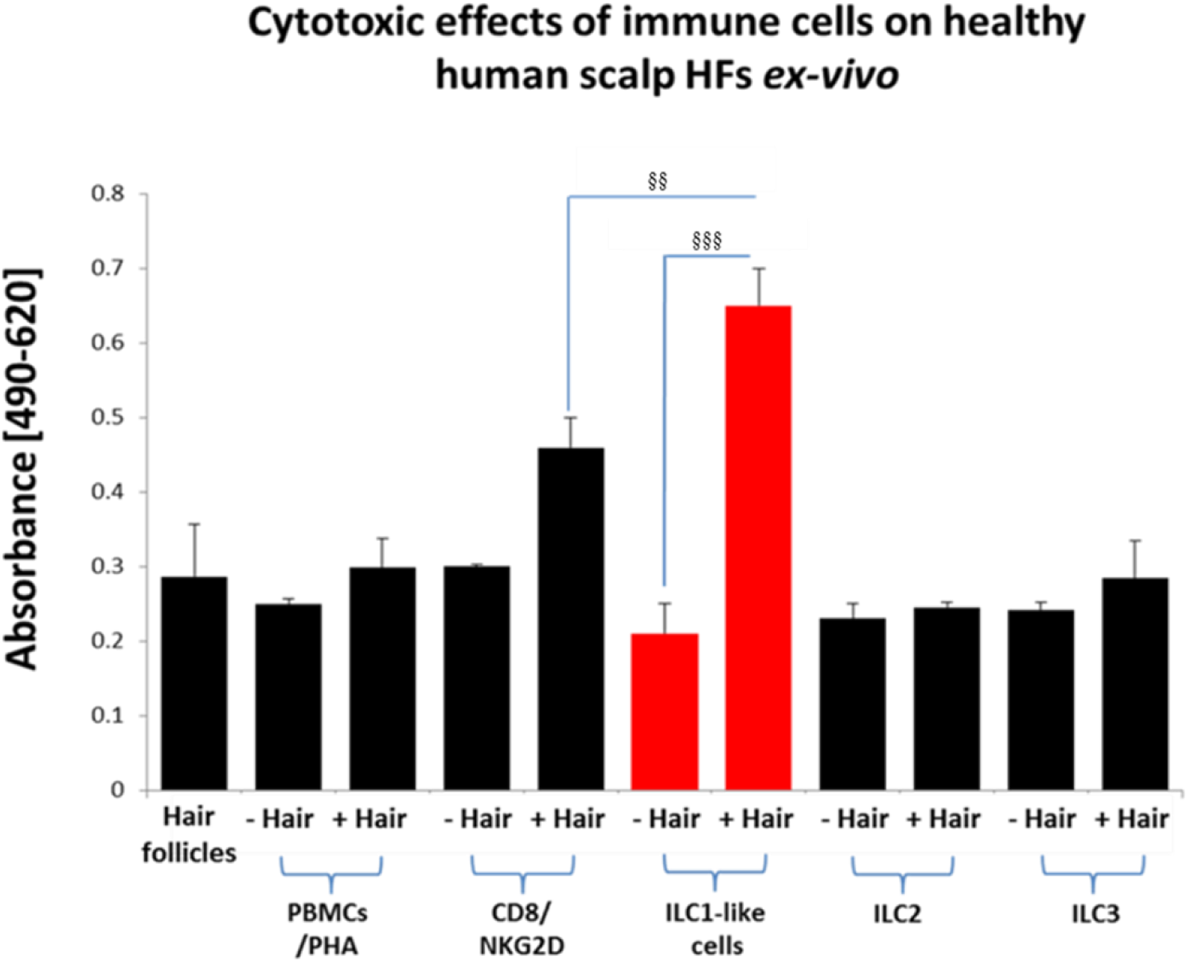
Cytotoxic effects of CD8+/NKG2D+ and ILC1-like cells on normal human scalp HF *ex vivo*. These cell populations were placed separately into wells with (+Hair) dissected HFs and without (-Hair). Cytotoxic effects of these cell populations on normal human scalp HF *ex vivo* was studied by measuring the spontaneous release of lactate dehydrogenase (LDH) from the microdissected HFs. Increased cytotoxicity of ILC1-like cells co-cultured with HFs compared to CD8+/NKG2D+, as well as to ILC2s and ILC3s and PBMCs/PHA cells. N=20-24 HFs/group derived from 3 independent donors analyzed in 3 independent HF organ culture experiments. Following Shapiro-Wilk test and Dunn’s test ^§^*p* < 0.05, ^§§^*p* < 0.01, ^§§§^*p* < 0.001.

**Figure. 4.**
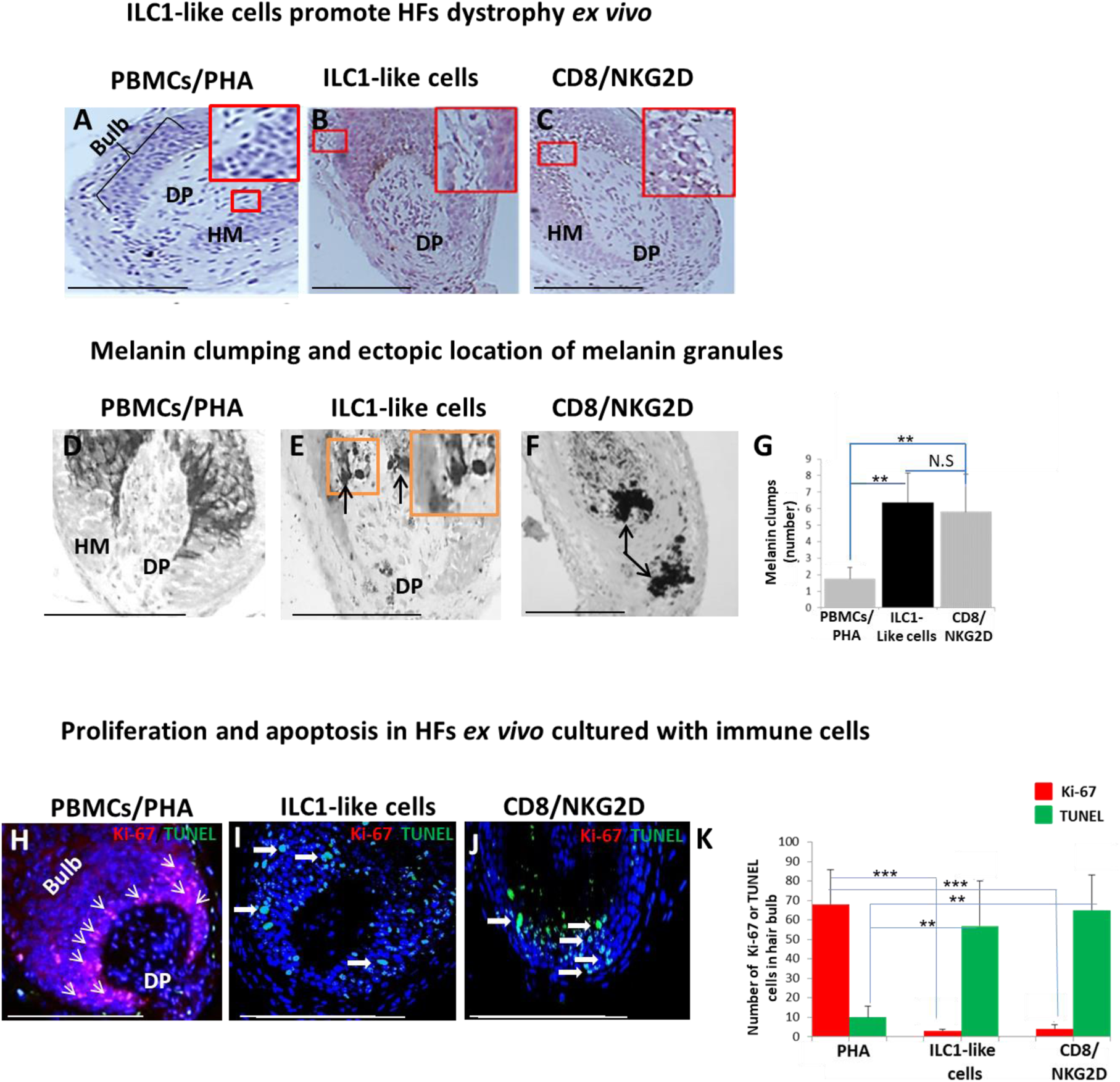
HFs dystrophy, melanin clumping and apoptosis in normal human scalp HF *ex vivo* co-cultured with ILC1-like and CD8+/NKG2D+ cells. (**A-C**) H&E staining revealed undifferentiated and prominent matrix cells, condensed dermal papilla and the appearance of apoptotic cells, N=15-19 HFs/group from 3 independent donors. (**D-G**) Masson-Fontana histochemistry revealed melanin clumping and ectopic location of melanin granules only in HFs co-cultured with CD8+/NKG2D+ and ILC1-like cells, but not in HFs cultured with PBMCs/PHA. N=7-11, HFs/group from 3 independent donors. Following Shapiro-Wilk test, Student’s *t*-test: \**p* < 0.05, \*\**p* < 0.01, \*\*\**p* < 0.001. (**H-K**) HFs co-cultured with ILC1-like or CD8+/NKG2D+ cells showed a significantly decreased proliferation (pink, arrowhead) and increased apoptosis (green, wide arrows). N=6 HFs/group from 2 independent donors, 3 areas were evaluated per section. Following Shapiro-Wilk test, Student’s *t*-test: \**p* < 0.05, \*\**p* < 0.01, \*\*\**p* < 0.001 in the anagen hair bulb compared to HFs cultured with PBMCs/PHA. Scale bars, 50 µm. **DP** - dermal papilla, **HM** - hair matrix.

### ILC1-like cells induce HF immune privilege collapse *ex vivo* via NKG2D stimulation

Given that AA cannot occur without the prior collapse of HF immune privilege [HF-IP] (Gilhar et al., 2012; Bertolini et al., 2020), we also investigated the impact of ILC1-like cells on key HF-IP markers. Indeed, the co-culture of HFs with ILC1-like cells triggered IP collapse, as evidenced by ectopic and overexpressed HLA-A,B,C, ß2-microglobulin (ß2 MG), and HLA-DR, along with overexpression of the “danger”/tissue distress signals, MICA and CD1d, which interact with and stimulate NKG2D (Uchida et al., 2021; Fan et al., 2022) as compared to HFs interacting with PBMC/PHA or with ILC3 cells (**Figure 5A, B, C, D, E and F**).

**Figure. 5.**
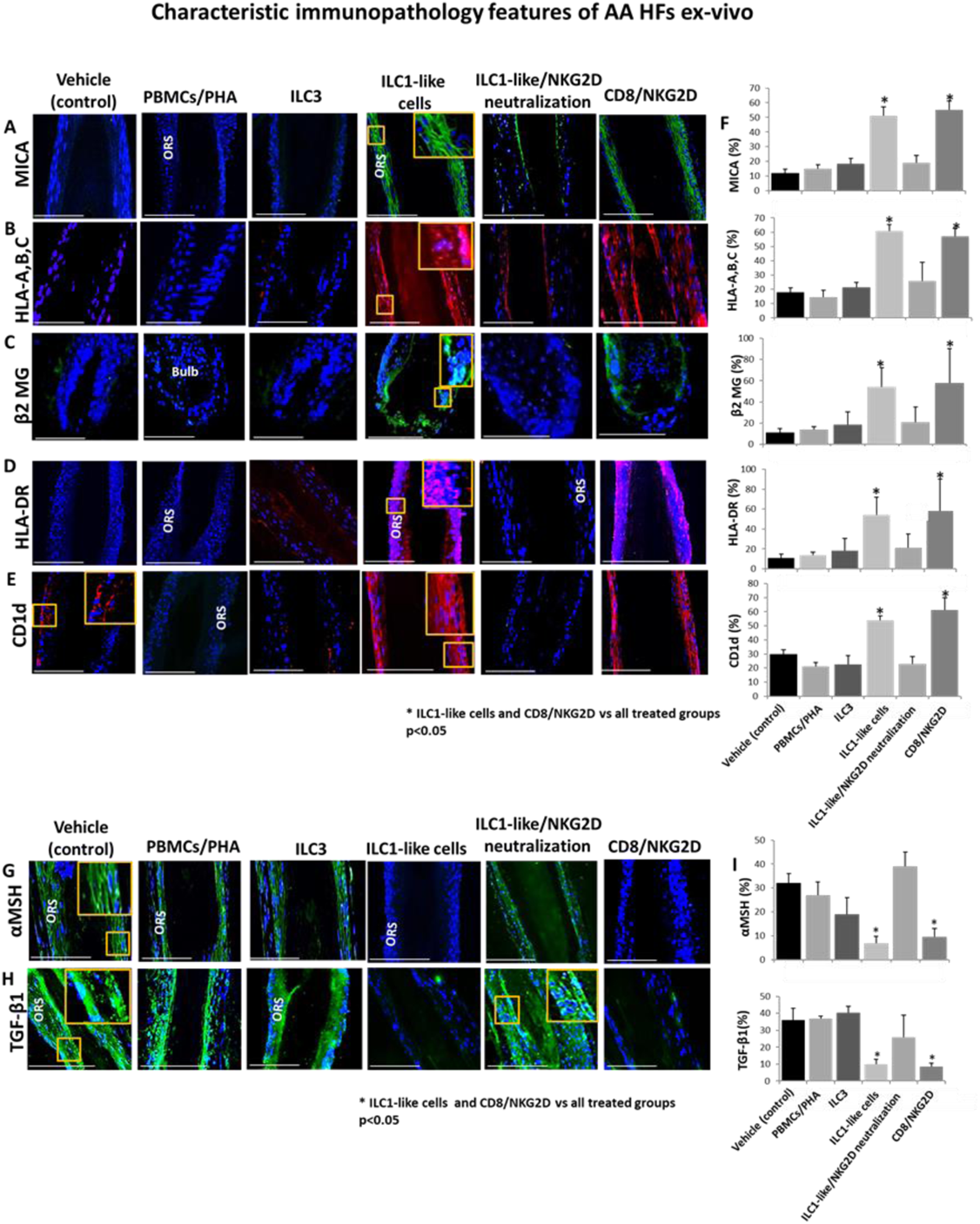
Characteristic immunopathology features of AA HFs. (**A**) MICA, (**B**) HLA-A,B,C, (**C**) β2 MG, (**D**) HLA-DR, and (**E**) CD1d, expression by HFs epithelium, which had been co-cultured with either ILC1-like or CD8+/NKG2D+ cells but not in the control HFs, which had been co-cultured with either ILC3s, PBMCs/PHA, ILC1-like/NKG2D neutralization or in the untreated HFs. (**F**) quantitation. (**G**) The immune inhibitory HF immune privilege guardians, α-MSH and (**H**) TGF-β1 almost disappeared in HFs/ILC1-like cells and HFs/NKG2D but were prominently present in ILC1-like/NKG2D neutralization and control HFs, N=9-12 HFs/group from 3 independent donors, 3 areas were evaluated per section. Following Shapiro-Wilk test, Student’s t-test, \**p* < 0.05. Scale bar, 100 µm. **ORS** - outer root sheet.

Notably, quantitative immunohistomorphoemtry (qIHM) also showed that protein expression of the immunoinhibitory HF-IP guardians, TGF-β1 and α-MSH (Gilhar et al., 2012; Bertolini et al., 2020; Paus et al., 2018; Ito et al., 2004), almost disappeared in the epithelium of HFs co-cultured with autologous ILC1-like or CD8+/NKG2D+ cells (=positive control) (**Figure 5G, H and I**), while these critical HF-IP guardians were still prominently expressed in negative control HFs (**Figure 5G, H and I**). Importantly, adding anti-NKG2D blocking antibodies prevented HFs IP collapse and preserved the IP in the ILC1-like/NKG2D treated group (**Figure 5 G, H and I**).

This demonstrates that autologous ILC1-like cells induce human HF-IP collapse *ex vivo* – incidentally, the first time that the induction of IP collapse by ILC1-like cells has been documented in an intact human tissue/organ.

### ILC1-like cells are activated by “stressed” HFs and induce premature catagen development via IFN-γ secretion

Next, we examined how autologous ILC1-like cells impacted on human HF cycling, given that premature induction of apoptosis-driven HF regression (catagen) is one of the hallmarks of AA (Gilhar et al., 2012; Bertolini et al., 2020; Messenger et al.,1986). This showed that ILC1-like cells significantly accelerated the transformation of anagen into catagen HFs *ex vivo* (Paus et al., 2005) compared to all three negative controls (ILC2, ILC3 or PBMCs/PHA) – thus eliciting the third hallmark of the AA phenotype besides HF-IP collapse and dystrophy *ex vivo* (Gilhar et al., 2012; Bertolini et al., 2020; Messenger et al., 1986) (**Figure 6A**), just as we had previously shown for Vδ1+ γδT cells (Uchida et al., 2021). As expected (Gilhar et al., 2012; Pratt et al., 2017; Bertolini et al., 2020; de Jong et al., 2018; Xing et al., 2014), premature catagen induction was also seen with CD8+/NKG2D+ cells (=positive control), but not with any of the negative control cell populations (**Figure 6A**).

**Figure. 6.**
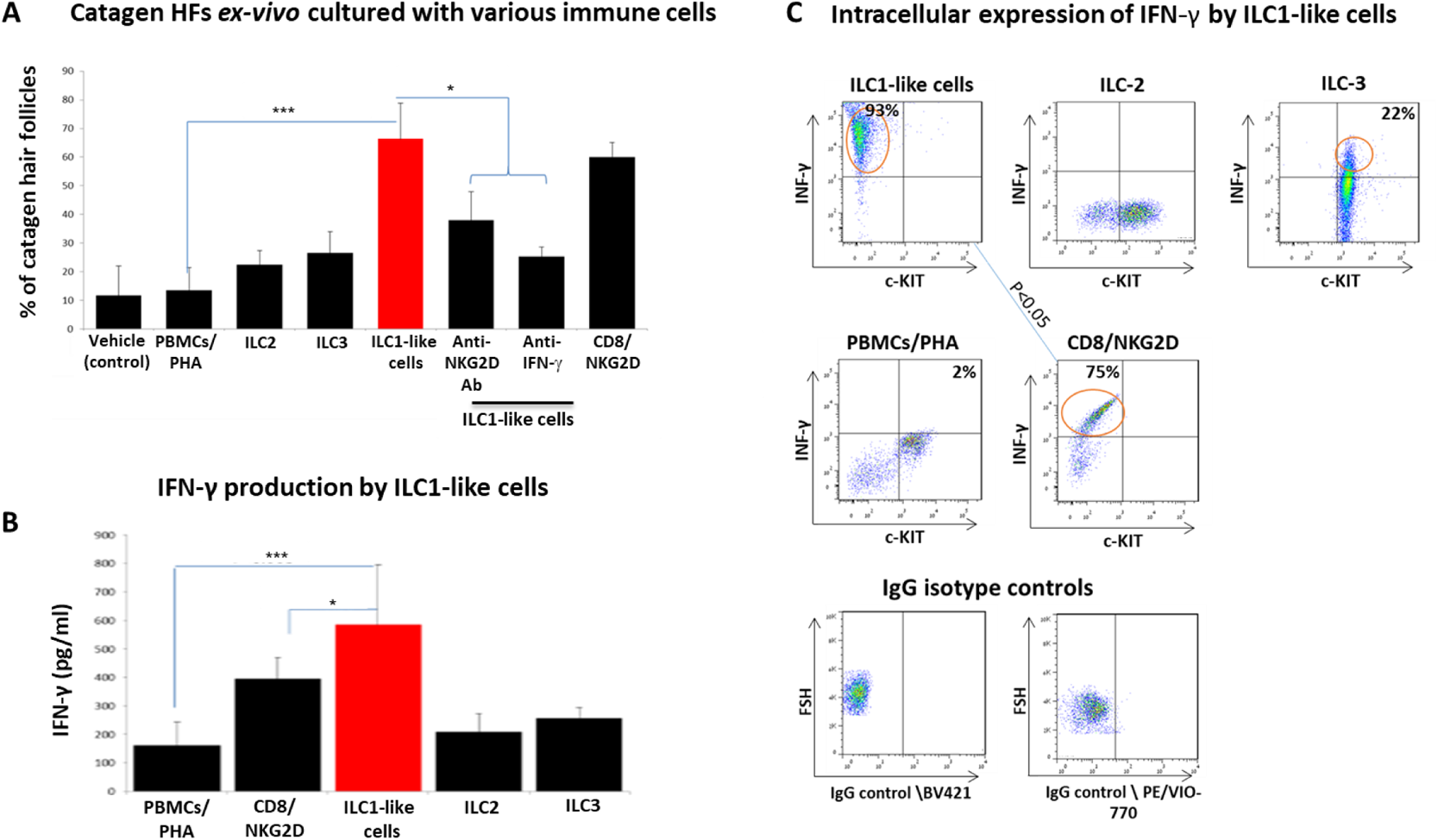
Transition of anagen to catagen HFs following culture with ILC1-like or CD8+/NKG2D+ cells in human scalp HF *ex vivo*. (**A**) These immune cells significantly accelerated the transformation of anagen HFs into catagen HFs *ex vivo* compared to ILC2, ILC3, PBMCs/PHA and neutralizing anti-IFN-γ, anti-NKG2D antibodies. N=28-34 HFs/group taken from 6 independent donors, Student’s *t*-test: \**p* < 0.05, \*\**p* < 0.01, \*\*\**p* < 0.001. (**B**) ELISA analysis revealed increased IFN-γ production by ILC1-like cells/HFs compared to production by CD8+/NKG2D+ cells, ILC2s, ILC3s and PBMCs/PHA. N=6 healthy donors, 6×10^6^ cells from each donor. Following Shapiro-Wilk test, Student’s t-test: \**p* < 0.05, \*\**p* < 0.01, \*\*\**p* < 0.001. (**C**) FACS analysis revealed a significant increased intracellular IFN-γ expression in ILC1-like cells co-cultured with HFs compared to the effector CD8+/NKG2D+ and to ILC2s and ILC3s, N=6 blood donors, 1.5×10^6^ cells/blood donor. Student’s *t*-test, *p*<0.05.

ILC1-like cells prominently secrete IFN-γ (Seillet et al., 2021), i.e. the cytokine that we had shown to induce HF damage (dystrophy), premature catagen and HF-IP collapse most potently (Ito et al., 2004; Ito et al., 2005a). Therefore, we next investigated IFN-γ release in these co-culture experiments. ELISA analysis revealed that ILC1-like cells produced and secreted higher amounts of IFN-γ into the medium than all other cells co-cultured with “stressed” HFs, including CD8+/NKG2D+ cells (p<0.05) (**Figure 6B**). This suggests that ILC1-like cells possess even stronger HF cytotoxicity-, IP collapse- and dystrophy-inducing properties than CD8+ T cells, the classical effector cells of AAA (Gilhar et al., 2012; Pratt et al., 2017; Bertolini et al., 2020).

FACS analysis showed ILC1-like cell activation when these were co-cultured with “stressed”, autologous HFs (600 cells/HF), as evidenced by significantly increased intracellular IFN-γ expression by ILC1-like cells (Bernink et al., 2017) (93±11%) compared to the positive control (CD8+/NKG2D+, 75±9%, p<0.05) (**Figure 6C**) and the negative controls ILC2s, 11±1%, p<0.05; ILC3s, 28±5%, p<0.05; PBMCs/PHA, 2±1.3%, p<0.001) (**Figure 6C**).

When neutralizing anti-IFN-γ antibodies were administered into the medium of the organ culture, premature catagen development of HFs co-cultured with ILC1-like cells was significantly reduced (**Figure 6A**), strongly suggesting that premature catagen induction by ILC1-like cells depends on their IFN-γ secretion (Seillet et al., 2021). Importantly, reduced catagen induction was also seen after adding NKG2D-blocking antibodies to the medium (**Figure 6A**). This suggests that ILC1-like cell activation and IFN-γ secretion is induced by NKG2D-stimulating danger signals overexpressed by stressed HF epithelium, such as MICA. These findings further support the recognized central role of both IFN-γ and NKG2D in the initial stages of AA pathobiology (Gilhar et al., 2012; Paus et al., 2018; Ito et al., 2008; de Jong et al., 2018).

### ILC1-like cells suffice to induce AA lesions in healthy human scalp skin *in vivo*

Taken together, these clinically relevant *ex vivo* experiments documented that ILC1-like cells can indeed induce the hallmarks of AA in healthy human scalp HFs *ex vivo*: HF-IP collapse, HF dystrophy and premature catagen development (Gilhar et al., 2012; Paus et al., 2018). Therefore, we finally probed the hypothesis that ILC1-like cells may also suffice to induce human AA-like hair loss lesions *in vivo* using our established humanized AA mouse model (Gilhar et al., 2013a; Ghraieb et al., 2018; Gilhar et al., 2016). We had previously demonstrated that a macroscopic and histological phenocopy of human AA lesions can be rapidly induced experimentally in healthy human scalp skin xenotransplants on SCID/beige mice in vivo by the intradermal injection of enriched CD8/NKG2D, which are defined as PBMCs that have been cultured for 14 days in high-dose IL-2 (100 U/ml) according to our previously published characterization (Ghraieb et al., 2018; Gilhar et al., 2013a; Gilhar et al., 2013b; Gilhar et al., 2016; Keren et al., 2015). These cells are derived from healthy donors, i.e. in the absence of a specific genetic or autoimmune constellation.

For this, 10 SCID/beige mice were each xenotransplanted with three full-thickness human scalp skin grafts (3 mm) obtained from parietal skin regions of four healthy donors without a prior history of AA (males aged 37±6). Eighty-nine days after transplantation, i.e., when hair regrowth had occurred in all xenotransplants, the mice were randomly divided into three groups, and each mouse from each group received one intradermal injection of either autologous IL-12/IL-18/IL-33-preactivated ILC1-like cells (test), PBMCs co-cultured with a nonspecific mitogen (PHA; negative control), or enriched CD8/NKG2D cells (positive control). When measured 45 days later, significant AA-like hair loss was observed macroscopically in the xenotransplants injected with ILC1-like cells compared to the negative control, at about the same level of positive control xenotransplants (**Figure 7A**).

**Figure. 7.**
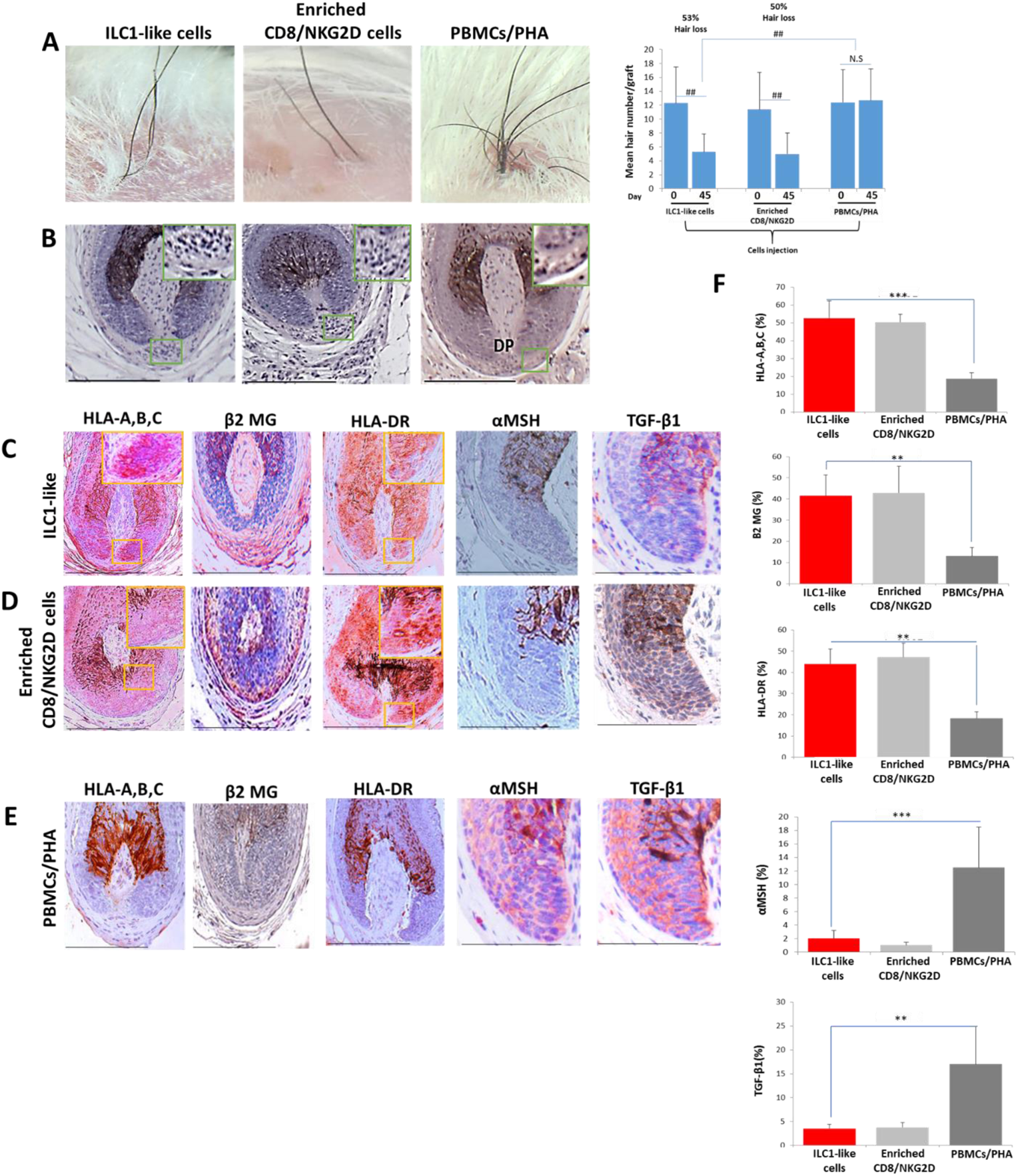
Development of AA in the humanized mouse model treated with ILC1-like cells. (**A**) Significant hair loss is observed following the injection of ILC1-like and enriched CD8/NKG2D cells, while in the PBMCs/PHA treated group, hair number remains almost constant. N=7-9 xenotransplants/group from 3 independent donors, Following Shapiro-Wilk test, Mann-Whitney *U* test: ^#^*p* < 0.05, ^##^*p* < 0.01. (**B**) HF dystrophy and perifollicular lymphocytic infiltrate around anagen HFs (H&E staining) combined with strong expression of (**C**) HLA-A,B,C, β2 MG, HLA-DR and downregulation of α-MSH and TGF-β1 in the ILC1-like cells and in (**D**) enriched CD8/NKG2D cells versus xenotransplants treated with (**E**) PBMCs/PHA (IHC staining) (**F**) quantitative data. N=5-9 xenotransplats/group from 3 independent donors. 4-5 defined reference areas were evaluated per section, and 3 sections per xenotransplants. Following Shapiro-Wilk test, Student’s *t*-test: \**p* < 0.05, \*\**p* < 0.01, \*\*\**p* < 0.001. Scale bar, 50 µm. **DP** - dermal papilla, **HM** - hair matrix.

To exclude that the above phenomena were not caused by residual human T-cells present in the transplants, an additional eight xenotransplanted mice were also injected intradermally once daily for 45 days with anti-CD3 antibodies (OKT3), in addition to injecting either ILC1-like or enriched CD8/NKG2D cells as described above (four mice each). This showed that anti-CD3 failed to abrogate hair loss induction in the mice treated with ILC1-like cells alone, but suppressed hair loss in the group treated with enriched CD8/NKG2D cells, as expected (**Figure 7----figure supplement 2A**). These findings invalidate the residual T-cell hypothesis.

### ILC1-like cells induce the characteristic immunopathology of human AA lesions *in vivo*

Immunohistology revealed that ILC1-like cells, just like autologous enriched CD8/NKG2D cells, induced a phenocopy of AA immunopathology in previously healthy human scalp skin *in vivo,* in sharp contrast to the negative control PBMCs/PHA group: HFs dystrophy, miniaturization and perifollicular lymphocytic infiltrate around anagen HFs (**Figure 7B**) as well as induction of HF-IP collapse (significantly increased expression of HLA-A,B,C, β2 MG and HLA-DR of the HF epithelium, along with downregulation of the immune privilege guardians, α-MSH and TGF-β1) (**Figure 7C and D**). In contrast, negative control xenotransplants injected with PBMCs/PHA showed normal anagen HFs and a significantly lower expression of HLA-A,B,C, β2 MG and HLA-DR, paired with the expected normal expression levels α-MSH and TGF-β1 protein (**Figure 7E**) as assessed by qIHM (**Figure 7F**). In addition, histology and quantitative immunohistomorphometry confirmed the preventive effect of anti-CD3 antibodies in inducing an AA-like phenotype in xenotransplants treated with enriched CD8/NKG2D cells, but not in those treated with ILC1-like cells (**Figure 7----figure supplement 2B, C, D, E, F and G and Figure 7----figure supplement 3**).

In line with the key role of IFN-γ in the development of AA (Gilhar et al., 2012; Gilhar et al., 2019a), IFN-γ + cells were found to be increased around the bulb of xenotransplants injected with ILC1-like cells, even in the presence of the anti-CD3 antibody (OKT3), or with enriched CD8/NKG2D cells, but not with PHA-treated PBMCs or enriched CD8/NKG2D cells in the presence of OKT3 (**Figure 7----figure supplement 2A, B, C, D, E, F and G and Figure 7----figure supplement 3A, B, C and D**).

### ILC1-like cells in the experimentally induced AA lesions

Given that both, enriched CD8/NKG2D cells, and ILC1-like cells produce high amounts of IFN-γ, we then investigated the subtype of these cells around the bulb of control and treated xenotransplants, along with the frequencies of CD4+ T-cells. In enriched CD8/NKG2D cells injected xenotransplants, the number of CD8+ cells and CD4+ T-cells was significantly increased as compared to ILC1-like cells (p<0.001,p<0.001) (**Figure 7----figure supplement 4A, B, C and D**) while dense infiltration of ILC1-like cells was found only in xenotransplants treated with the purified ILC1-like cells (**Figure 7----figure supplement 4E and F**).

Interestingly, qIHM also showed that the peri- and intrafollicular distribution and mean number of ILC1-like cells in human skin xenotransplants injected with enriched CD8/NKG2D cells imitated that of ILC1-like cells seen in spontaneously developed hair loss lesions of AA patients, further supporting the role of ILC1-like cells in human AA (**Figure 1A, B, C and D, Figure 8A, B, C and D and Figure 8----figure supplement 1B**). Yet, CD8+/NKG2D+ lymphocytes significantly outnumbered ILC1-like cells in the experimentally induced AA lesions (p<0.01) (**Figure 8E and F**), just as they do in human AA patients (**Figure 1 E and F**).

**Figure. 8.**
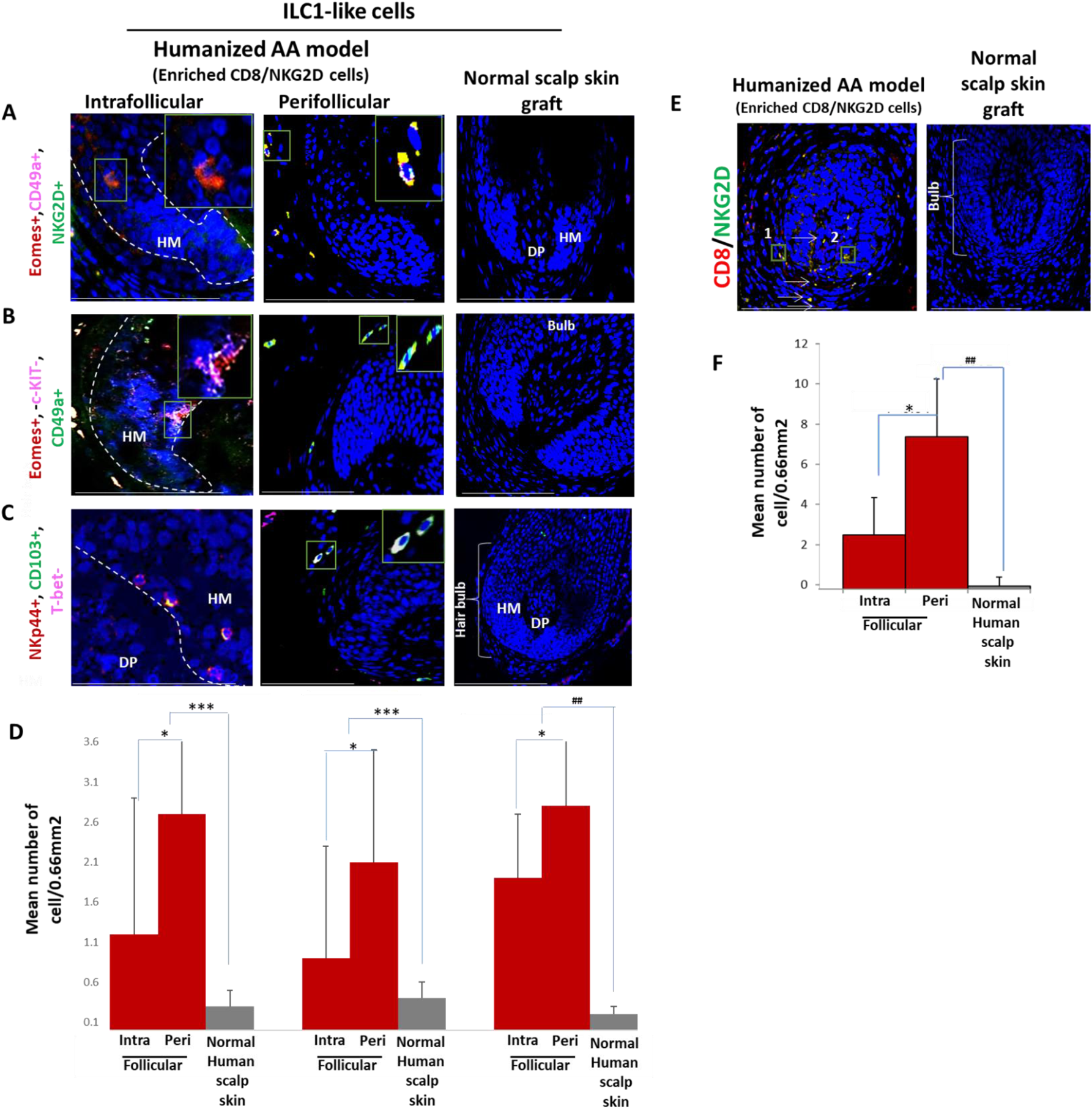
Immunofluorescence microscopy analysis of ILC1-like and enriched CD8/NKG2D cells in AA-induced xenotransplant. **(A)** EOMES+, CD49a+ and NKG2D+ around HF in normal scalp skin, intrafollicular and perifollicular ILC1-like cells infiltrates in AA-induced xenotransplants (**B**) EOMES+,c-KIT−,CD49a+ and (**C**) NKp44+, CD103+, T-bet-ILC1-like cells. Absence of these cells in normal scalp xenotransplant. (**D**) Quantitation. (**E**) CD8+/NKG2D+ cells around HF in AA-induced xenotransplant versus absence of the cells in normal xenotransplant. (**F**) The quantitative data demonstrate the significant increased CD8+/NKG2D+ cells in HFs of AA humanized mice compared to normal scalp xenotransplants. N=6 xenotransplants/ group from 3 independent donors, 3 areas were evaluated per section. Following Shapiro-Wilk test, Student’s *t*-test: \**p* < 0.05, \*\**p* < 0.01, \*\*\**p* < 0.001. Mann Whitney *U* test: ^#^*p* < 0.05, ^##^*p* < 0.01. Scale bar, 50 µm. **DP** - dermal papilla, **HM** - hair matrix, White arrow-c-KIT stained melanocyte.

## Discussion

The current study is the first to phenotypically and functionally explore the role of ILC1-like cells in human AA *in vivo* and *ex vivo*. Here, we show that ILC1-like cells are increased in AA lesions provide the first functional evidence that expanded circulating autologous human ILC1-like cells suffice to induce all hallmarks of the AA hair loss phenotype (premature catagen, HF dystrophy and HF-IP collapse) in previously healthy, organ-cultured human scalp HFs *ex vivo* and in human scalp skin xenotransplants *in vivo,* where they also cause the characteristic clinical hair loss phenomenon. This also provides the first unequivocal functional evidence of a key role of ILC1-like innate lymphocytes in a model human autoimmune disease, and thus identifies these lymphocytes as important novel targets in future AA therapy.

Mechanistically, we demonstrate that IFN-γ secretion and NKG2D signaling are both required for this AA-pattern HF damage to occur (**Figure 9**). That ILC1-like cells alone can induce all hallmarks of AA in a healthy human (mini-)organ *ex vivo* and *in vivo*, presumably in an autoantigen-independent manner, also demonstrates that these innate/transitional lymphocytes interact directly with human HFs, rather than affecting them only indirectly. We also show that resident T-cells in human scalp skin transplants are not responsible for the AA-inducing effects of ILC1-like cells *in vivo* (**Figure 7----figure supplement 2 and Figure 7----figure supplement 3**).

**Figure. 9.**
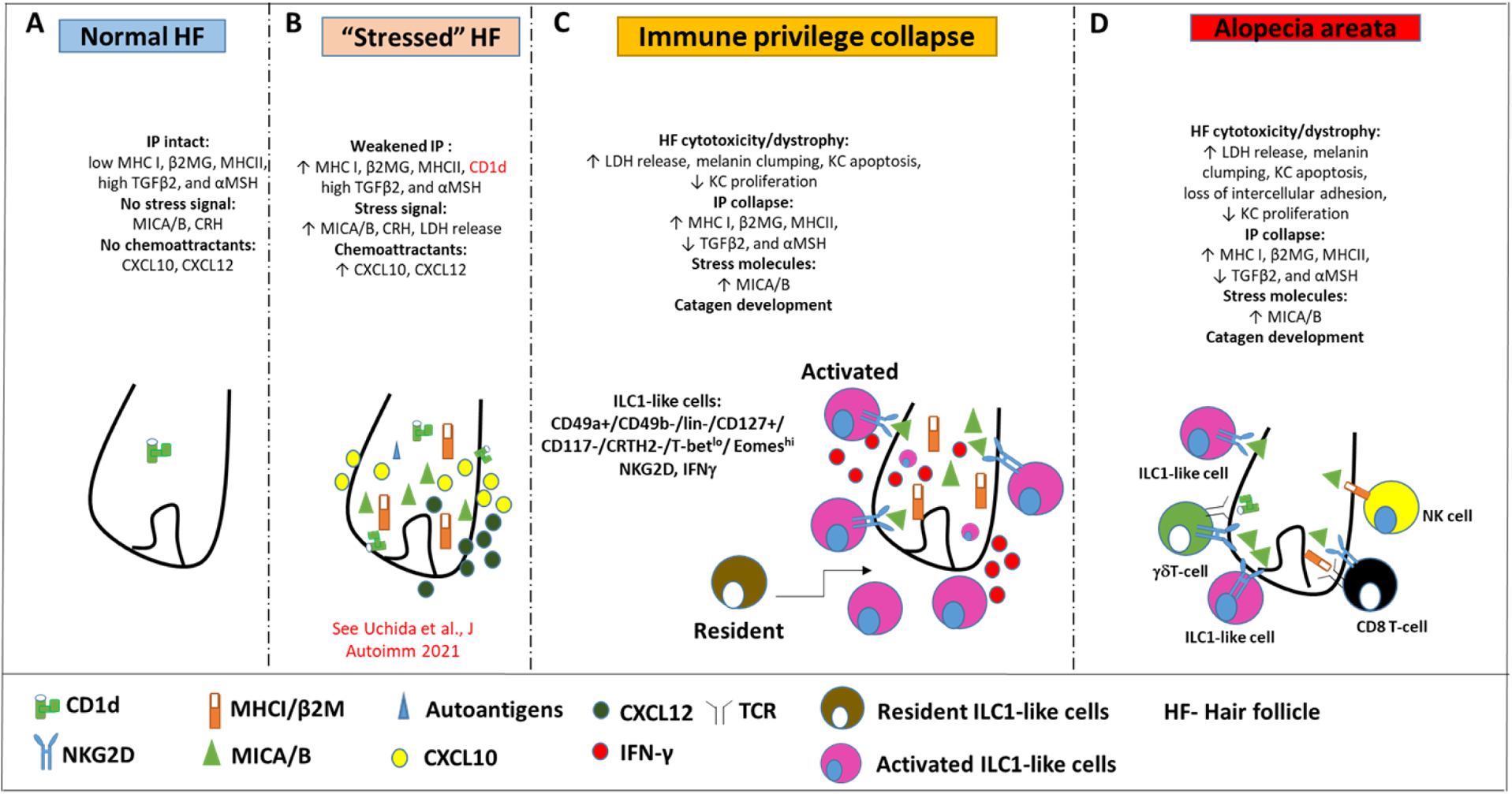
Pathobiology scenario: How ILC1-like cells can induce AA. (**A**) ILC1-like cells are rarely detected around the bulb of healthy human scalp HFs, which exhibit relative immune privilege and low or absent expression of MICA and MHC class I, and CD1d. (**B**) Various tissue stressors (in the current study: hair follicle microdissection and organ culture), can transiently weaken the hair follicle’s physiological immune privilege by upregulating the expression of MHC class I, MICA (a key activating NKG2D ligand), and of CD1d, along with the secretion of chemoattractants such as CXCL12. (**C**) This recruits and activates ILC1-like cells, which migrate towards the “stressed” hair follicle and secrete IFN-γ, thus ultimately inducing HF-IP collapse. (**D**) Either alone or in conjunction with other recognized AA-inducing immune cells (i.e., CD8+ T cells which recognize hair follicle autoantigens now exposed by ectopically expressed MHC class I; NK cells, and *γδ*TCs), ILC1-like cells can then induce the full AA phenotype, characterized by HF-IP collapse, premature hair follicle regression (catagen), and hair follicle dystrophy.

Our study demonstrates that CD8^+^ T cells, which have long been thought to represent the central players in AA pathobiology (Pratt et al., 2017; Xing et al., 2014; de Jong et al., 2018; Gilhar et al., 1998; Wang et al., 2016; Paus et al., 1993), are not the only drivers of disease (Gilhar et al., 2019a; Pratt et al., 2017; de Jong et al., 2018; Paus et al., 1993), and are no joined not only by NK cells (Ito et al., 2008; Gilhar et al., 2013a) and γδT cells (Uchida et al., 2020; Uchida et al., 2021), but also by ILC1-like lymphocytes. The study further supports the concept that the characteristic hair loss pattern we diagnose as AA phenotype, does not *always* represent a classical, autoantigen- and autoreactive CD8+/NKGD2+ T-cell-dependent autoimmune disease (“autoimmune AA” [AAA]), but can also reflect non-autoimmune pathomechanisms that may perhaps best be defined as non-autoimmune AA subtype (NAIAA) (Bertolini et al., 2020; Paus et al., 2018) (**Figure 9**). In a state of prolonged HF-IP collapse and thus chronic exposure of HF-associated autoantigens to (pre-existent?) autoreactive CD8+ T cells, it is well conceivable that an AA-subtype that began as NAIAA can over time transform into the AAA-variant, thus explaining the chronic-intermittent course that is seen in so many autoimmune diseases.

The novel concept of ILC1-induced NAIAA mandates a differential, personalized management approach to future AA therapy, which tailors treatment to the specific pathobiology at hand in any given AA patient. This must now include the targeting of pathogenic, potently IFN-γ-secreting ILC1-like cells, at least when these are seen to be increased in lesional AA biopsies.

Although the number of ILC1-like cells in lesional AA skin was significantly lower than that of CD8+ cells (**Figure 1**) this does not rule out a crucial role of the cells in spontaneous AA development in patients. In fact, we demonstrate here that ILC1-like cells are even more potent IFN-γ producers than CD8+ T-cells. Also, despite the relatively low numbers of ILC1-like cells, their selective tissue distribution makes them ideally localized to provide an early source of cytokines to initiate/trigger pro-inflammatory immune responses directed against distressed tissues (Vivier, 2021), as documented here in our co-culture assay with “stressed” human scalp HFs *ex vivo*. That IL-12 and IL-18 were among the cytokines used here to facilitate ILC1-like cell isolation from human PBMCs is also interesting in the context of our most recent observation that local IL-12 signaling, supported by IL-18, may be involved in the early stages of AA development by stimulating IFN-γ production from resident IL-12Rß2+ immune cells, eventually leading to HF-IP collapse (Edelkamp et al., 2021). In fact, treatment with IL-12+IL-18 of healthy HFs selectively enriches IFN-γ-inducible genes and promotes the release of IFN-γ into the medium and thus HF-IP collapse. These responses were abrogated by co-administration of a selective TYK2 inhibitor (Edelkamp et al., 2021), which confirms a key role of IL-12, whose receptor utilizes a TYK2 and Janus kinase 2 pair for downstream signal transduction (Ullrich et al., 2020; Krueger et al., 2022), These preliminary findings suggest that IL-12 is a key effector cytokine in promoting pathogenic IFN-γ secretion and HF-IP collapse. Given that ILC1 are IFN-γ producing cells (Nabekura et al., 2021b; Resende et al., 2017; Quintino-de-Carvalho et al., 2022), our data therefore encourage one to dissect, next, the exact role of IL-12/IL-12R-mediated signaling in activating, attracting and/or expanding ILC1-like cells and in stimulating their IFN-γ secretion in the early stages of human AA pathogenesis.

An immunopathology-initiating role of ILC1-like cells is not unique to AA and has also been proposed in other inflammatory conditions (Kim et al., 2021), including vitiligo (Tulic et al., 2019), which shares some pathogenesis features with AA (Harris, 2013a; Tomaszewska et al., 2020), inflammatory bowel disease (McDonald et al., 2018; Bernink et al., 2013; Tang et al., 2019), lupus erythematosus (Guo et al., 2019), and the aggravation of atherosclerosis (Wu et al., 2018). Yet, to the best of our knowledge, the current study is the first to demonstrate that these innate immunocytes can indeed induce the full disease-mimicking immunopathology phenotype in a previously healthy human (mini-)organ. Also, abnormalities in the crosstalk between ILC1-like cells and gut microbiota have been observed in various diseases (Jiao et al., 2020). Therefore, it is interesting to ask whether the microbial dysbiosis that has been reported in scalp skin of AA patients (Pinto et al., 2020) may activate the very few, strategically positioned perifollicular ILC1-like cells present in healthy human scalp skin through abnormal crosstalk with HF microbiota (Lousada et al., 2021). This hypothesis can now be explored using our *ex vivo* and humanized AA mouse model.

Collectively, our study introduces IFN-γ-secreting ILC1-like lymphocytes as important novel players in human AA pathobiology and identifies them as new therapeutic intervention targets. Their strategic location, their capability to recognize and respond to HF distress signals such as MICA and selected chemokines, the excessive production of IFN-γ by ILC1-like cells, and their direct, pathogenic effects on human HFs *ex vivo* documented here, all support that these cells play a hitherto unappreciated role in early AA pathogenesis. Moreover, our study demonstrates that autoreactive CD8+ T cells are not indispensable for AA induction and further supports that non-autoimmune AA variants (NAIAA) (Bertolini et al., 2020) exist and that innate/transitional immune cells play an important role in AA pathobiology.

## Materials and Methods

### Patients, tissue and blood samples

For the *in situ* analyses we used archival paraffin-embedded biopsy specimens of human AA scalp skin lesions from the Department of Pathology, Rambam Medical Center (4 females, 12-35 years, mean age 20.5±9.5; 6 males, 6-38 years, mean age 18±12). Three of these AA patients showed active hair loss of the AA universalis phenotype while the other patients showed stable hair loss patches of the multifocal AA phenotype (Gilhar et al., 2012). One ten-year-old male patient had a positive family history of allergic rhinitis. AA was diagnosed both clinically and by histopathology, and none of the enrolled patients showed clinical evidence or had a personal history of other AA-associated autoimmune diseases (Gilhar et al., 2012; Meah et al., 2021).

Clinically healthy human skin scalp specimens were obtained from healthy volunteers undergoing cosmetic facelift surgery (3 females, 43-72 years, mean age 58±15; 5 males, 31-44 years, mean age 36±8).

The *ex vivo* experiments utilized scalp HFs from 20 healthy donors (13 males and 7 females) without a history of AA (31-63 years, mean age 49±12).

The *in vivo* experiments in the humanized AA mouse model (Ghraieb et al., 2018) used scalp skin pieces obtained from five healthy donors (4 males and 1 female, 34-45 years, mean age 38±5). For the *ex vivo* and *in vivo* experiments, frontotemporal human scalp skin specimens were obtained during elective cosmetic facelift procedures performed under general anesthesia, and 20 ml of autologous venous blood were drawn, both with informed written patient consent.

The study for both *ex vivo* and *in vivo* experiments was approved by the Institutional Ethics Committee of the Rambam Health Care Campus, Haifa, Israel (RMB-0182-14).

**For the *ex vivo* experiments**, frozen HFs sections were dehydrated for 40 minutes and incubated with acetone for 10 minutes at −20°C for fixation. Slides were dehydrated for 20 minutes and transferred to double-distilled water following three times wash with 1xphosphate buffered saline (PBS) (pH=7.4). Sections were blocked with suitable serum (horse/goat) for 30 minutes to prevent nonspecific binding and incubated at 4°C with primary Ab overnight. Slides were incubated with an appropriate biotinylated secondary Ab (FITC-conjugated goat anti-mouse Ab, Rhodamine-conjugated goat anti-mouse IgG Ab or Alexa Fluor 488-conjugated goat anti-rabbit IgG Ab) for 30 minutes, following three times wash with PBS.

**For the *in vivo* experiment**, five-micrometer paraffin sections were used. Antigen retrieval was for 20 min at 90°C in a microwave. Specimens were blocked for 30 min to prevent nonspecific binding and incubated with the first antibody (Ab) overnight, followed by a wash and incubation with biotinylated 2^nd^ Ab (Jackson ImmunoResearch, West Grove, PA), and subsequent binding with horseradish peroxidase conjugated streptavidin. Markers were revealed with AEC (red) (Aminoethyl Carbazole Substrate kit). Sections were then mounted and analyzed under a light microscope.

### Histochemistry, immunohistology and quantitative immunohistomorphometry (qIHM)

Five-micrometer paraffin sections of lesional AA biopsies and human scalp skin xenotransplants were processed for histochemistry or immunohistology. The following primary antibodies were used: anti-CD8 (Cell Marque-108M-95), anti-CD4 (DAKO-M7310), anti-HLA-A,B,C (Abcam-70328), anti-HLA-DR (Abcam-20281), anti-IFN-γ (Abcam-25101), anti-α-MSH (LSBio-C25584), anti-beta2 microglobulin (Abcam-218230) and anti-TGF-β1 (Santa Cruz-52893) (Ghraieb et al., 2018; Laufer Britva et al., 2020; Keren et al., 2015).

Since there is no single, highly specific surface marker for ILC1 cells, triple immunostaining was performed with three sets of antibodies that are routinely used for the identification of human ILC1-like cells (Talayero et al., 2016; Hawke et al., 2020a,b; Cruz-Zárate et al., 2018): a) NKp44+ (Bioss-YEYS3W) (Talayero et al., 2016), CD103+ (eBioscience 1401038-82) (Hawke et al., 2020b) and T-bet- (Santa Cruz-H3112) (Cruz-Zárate et al., 2018); b) c-KIT− (DAKO-MA512944) (Nagasawa et al., 2019), CD49a+ (R&D systems-AF5676) (Colonna et al., 2018) and EOMES+ (ThermoFisher-14-4877-82); and c) CD49a+ (R&D systems), EOMES+ (ThermoFisher-14-4877-82) and NKG2D+ (Novus-5c6) (de Jong et al., 2018). Skin-infiltrating CD8+NKG2D+ T-cells were double-immunostained by NKG2D (Novus-5c6)/CD8 (Cell Marque-108M-95) (de Jong et al., 2018). For negative control, the primary antibody was replaced with non-specific IgG1 and IgG2 isotype control. Hematoxylin and eosin (H&E) staining was performed on cryo- or paraffin sections as previously described (Keren et al., 2018). For the *ex vivo* experiments, HFs cryosections were dehydrated for 40 minutes and fixed with acetone for 10 minutes at −20°C (Ghraieb et al., 2018; Laufer Britva et al., 2020; Keren et al., 2015).

The following primary antibodies were used for immunohistochemistry (IHC) or immunofluorescence microscopy (IF) of key HF immune privilege markers (Bertolini et al., 2020; Paus et al., 2005) anti-HLA-A,B,C (Abcam-70328)/ anti-HLA-DR (Abcam-20281)/anti-MICA(Santa Cruz-20931)/anti-CD1d (Abcam-11076)/anti-α-MSH (LSBio-C25584)/anti-beta2 microglobulin (Abcam-218230) and anti-TGF-β1(Santa Cruz-52893).

The immunoreactivity patterns were assessed in standardized, well-defined reference areas by quantitative immunohistomorphoemtry (qIHM) by experienced, blinded observers, following our standard protocols for evaluating human HF immunology read-outs (Bertolini et al., 2014; Bertolini et al., 2016; Christoph et al., 2000; Harries et al., 2013b; Hardman-Smart et al., 2020), counting at least three reference areas each on three non-consecutive sections, presented randomly to the blinded observer(s). Specifically, immunoreactive cells around and within the HFs were counted in an area of 0.66 mm^2^.

For HLA-A,B,C, HLA-DR, MICA, CD1d, α-MSH, beta2 microglobulin and TGF-β1 image analysis was performed using Image J software. Protein expression was measured by calculating the percentage of staining coverage within the analyzed area.

**Masson-Fontana staining** (Abcam) was performed as described by us (Laufer Britva et al., 2020; Purba et al., 2016). Briefly, five-micrometer paraffin sections were deparaffinized and hydrated in distilled water. Slides were placed in mixed ammoniacal silver solution in a 58-60°C water bath and allowed adequate time for the temperature to equilibrate. Slides were then placed in warmed ammoniacal silver solution for 30-60 minutes or until the tissue section became yellowish/brown in color. Counterstaining was performed with Nuclear Fast Red Solution for 5 minutes.

### TUNEL analysis

Apoptotic cells were evaluated using a commercial TUNEL kit (Roche) with antidigoxigenin fluorescein labeling and according to manufacturer’s protocol. Ki-67 (Invitrogen) was visualized using Alexa Flour 594-conjugated goat anti-mouse (Jackson, 115-585-062). Sections were counterstained by DAPI (Thermo Fisher Scientific). Staining was visualized using a confocal Microscope - Zeiss LSM 700. Quantification was performed as previously described (Peters et al., 2006).

### Immunohistology

Slides were photographed using immunofluorescence confocal microscopy and compared systematically by qIHM in standardized, defined tissue compartments. Mouse skin served as a negative control. Three non-consecutive sections were analyzed per patient.

### Isolation, characterization, and culture of circulating ILC1-like, ILC2, ILC3 and CD8+/NKG2D+ cells

ILC2 and ILC3 cells were used as negative controls, while CD8+/NKG2D+ cells were used as a positive control to evaluate the ILC1-like cells cytotoxic effects on HFs. The cells were cultured and induced to expand, as we have previously described (Keren et al., 2018). The isolation and characterization of ILC1-like, ILC2 and ILC3 cells by FACS cell sorting or MACS was performed as we previously described (Keren et al., 2018; Mjösberg et al., 2011; Mora-Velandia et al., 2017; Creyns et al., 2020).

Autologous human PBMCs were isolated from heparinized whole blood from healthy donors by Lymphoprep density gradient centrifugation (Alere Technologies, Norway). Cells were frozen for further assays (70% FBS, 20% RPMI1640 and 10% DMSO) or cultured at a seeding density of 3×10^6^ cells/ml in 24 wells plate with medium (RPMI 1640, 10% human AB serum, 1% penicillin-streptomycin antibiotics, 2 mML glutamine) in the presence of different cytokines for cells expansion.

The following components required for the expansion of various immune cell populations:

ILC1-like cells (Silver et al., 2016): IL-18(1 µg/1 ml) (CYT-269(A), IL-33((1.5 µg/5 ml) (BLG-581802), IL-12 (1.5 µg/5 ml) (BLG-573002).

ILC2 (Creyns et al., 2020): IL-7 (10 ng/ ml) (BLG-581904), IL-25 (100 ng/ml) (C792-50), IL-2 (50 ng/m) (Prospec-Cyt-209-b).

ILC3 (Keren et al., 2018): AHR (200 nM) (BML-GR2060100), IL-2 (100 U/mL). CD8+NKG2D+ (Gilhar et al., 2013a): IL-2 (100 U/ml).

PHA (Gilhar et al., 2013a): PHA (10 µg/ml) (Sigma-C1668).

On days three and five, half of the medium was either frozen for further analysis or discarded and replaced with fresh medium containing cytokines. After seven days, cells were sorted by FACS Aria (FACSAria™ III Cell Sorter, BD Biosciences, USA) and in the case of ILC3 further enriched by MACS for negative selection of CD3+ cells (see **Figure 3----figure supplement 5**).

### Flow cytometry sorting for ILC1-like, ILC2 and CD8+/NKG2D+ cells

Cells were cultured for one week, collected and washed with PBS containing 1% BSA and 2% PSN. Surface cells were stained with antibodies to the PE-conjugated lineage cocktail that includes antibodies against CD1a (BLG-300105), CD3 (BLG-300-307), CD14 (BLG-367103), CD19 (BLG-302207), CD34 (BLG-343605), CD123 (BLG-306005), CD11c (BLG-301605), BDCA2 (BLG-354203), FcεR1α (BLG-334609), TCRαβ (BLG-306707), TCRγδ (BLG-331209), CD56 (BLG-362565) (Hawke et al., 2020b).

Following gating on lineage, cell population cells were sorted as follows:

**ILC1-like cells** – APC-conjugated-CD127+ (BLG-351315), PE/CY7-conjugated-CD161+ (BLG-339917), and FITC-conjugated-NKp44 + (SC-53597), Brilliant Violet 421TM anti-human CD117- (c-KIT) (BLG-313215) and APC/CY7-conjugated-CRTH2- (BLG-350113). (Santa Cruz H3112) (Talayero et al., 2016; Hawke et al., 2020a).

**ILC2 cells** – APC-conjugated-CD127+ (BLG-351315), PE/CY7-conjugated-CD161+ (BLG-339917), Brilliant Violet 421TM anti-human CD117+ (c-KIT) (BLG-313215) and APC/CY7-conjugated-CRTH2+ (BLG-350113) (Cruz-Zárate et al., 2018; Nagasawa et al., 2019).

**CD8+/NKG2D+ cells** – CD8 (Cell Marque-108M-95)/NKG2D(Novus-5c6) (Ito et al., 2008). The cells were sorted using a FACS Aria instrument with software (BD Biosciences). The sorted cells were collected in a tube with medium enriched with 20% human serum. Afterward, the cells were centrifuged, suspended, counted and co-cultured with HFs or used for ELISA assay.

Compensation was done using Comp-Beads (BDTM Biosciences) and data were analyzed using FlowJo software.

### Magnetic isolation of ILC3 subsets

Separation was performed using anti-CD3 antibodies conjugated to ferromagnetic microbeads (Miltenyi Biotec, Bergisch Gladbach, Germany) and directed through a cell separation column containing a magnetic field (Miltenyi Biotec). For the purification of ILC3s, CD3−sorted cells were collected and stained with anti-NKp44-PE conjugated to ferromagnetic microbeads (Miltenyi Biotec) and directed through a cell separation column containing a magnetic field (Keren et al. 2018).

Finally, cells were co-cultured with autologous HFs *ex vivo* (see below) or used for different assays (Keren et al., 2018).

### Co-culture of autologous ILC1-like cells with “stressed” human scalp hair follicles *ex vivo*

Experimental induction of HF-IP collapse by IFN-γ is the standard *ex vivo*-assay system for interrogating key elements of AA-related human HF immunopathology (Ito et al., 2004; Bertolini et al., 2016; Kinori et al., 2012). We have recently complemented this assay by co-culturing key immunocytes in AA pathogenesis (CD8+ T cells, γδTCs) directly with organ-cultured human scalp HFs *ex vivo* (Uchida et al., 2021).

For this, healthy human anagen scalp HFs were collected and microdissected as described (Ito et al., 2004), and HFs were placed individually into a 96-well plate slot with 100 µl supplemented medium (William’s E plus 1% penicillin-streptomycin antibiotics, 1% L-glutamine (Invitrogen-Gibco), 0.01% hydrocortisone (Sigma-Aldrich) and 0.01% insulin (Sigma-Aldrich) (Langan et al., 2015).

As we have documented in detail elsewhere (Uchida et al., 2021), on day 1 after initiation of organ culture, the HFs are markedly, but transiently stressed by the trauma of microdissection and the transfer to a harsh, hyperoxygenated *ex vivo* culture environment. This results in significantly increased LDH activity release and up-regulation of CXCL12 and CXCL10 expression as well as in a transient, partial weakening of the HFs physiological immune privilege. The latter was evidenced by increased protein expression of MHC class Ia, β2-microglobulin and MICA/B but no change in the expression of IP guardians such as αMSH and TGFβ2. As reported before, all these “HF stress” indicators normalize on day 3 of organ culture (**Table supplement 1**) (Taken from: Uchida et al., 2021). Thus, due to their expression of NKG2D ligands (MICA/B) CXCL12 and CXCL10 secretion, and transiently weakened HF immune privilege, the stressed (day 1) HFs can attract and interact with immune cells expressing NKG2D receptors and are primed to elicit anti-HF immune responses *ex vivo* (Uchida et al., 2021).

Therefore, organ culture-stressed day 1 HFs (1HF/well) were co-cultured in supplemented William’s E medium from day 1 until day 6 with one of 5 different immune cell populations: (1) ILC1-like cells (100 µl/600 cells per well), either alone or in combination with anti-IFN-γ antibody (10 µg/ml, R&D Systems, MAB285) or NKG2D neutralizing antibody (5 µg/ml, R&D Systems, MAB139-100); ILC1-like cells demonstrated cytotoxic effect on HFs with 600 cells per well, while (2) CD8/NKG2D cells demonstrated similar effect only with 100 µl/3500 cells per well. Therefore100 µl/3500 per well were used for the following control groups: (3) ILC2s cells; (4) ILC3s or (5) PHA cultured PBMCs.

The medium was not replaced in order to avoid losing any immunocytes. Basic HF biology read-out parameters were assessed by evaluating the Ki-67/TUNEL ratio, values of LDH release, HF pigmentation and hair shaft production *in situ*, all of which indicated that the HFs did not suffer major damage after 6 days of organ culture. At the end of the experimentation, the HFs were photo documented and cryopreserved in optimal cutting temperature (OCT) blocks. Cytokine release into the culture medium by ELISA was analyzed as previously described (Zook and Kee, 2016).

### Flow cytometry analysis for characterization of ILC1-like cells

PBMCs were isolated from healthy blood via centrifugation on ficol/Hypaque and cultured for seven days in a medium composed of RPMI 1640, 10% human AB serum, 1% L-glutamine and 1% PSN. The medium was changed as needed. Seven days later, six hours prior to FACS staining, cells were then collected (1-1.5×10^6^ cells/tube), centrifuged at 1200 RPM for 5 min and washed twice in staining buffer (1 ml of 1% Bovine Serum Albumin [BSA] in 1x sterile PBS). First antibodies (as described above, flow cytometry sorting) were used at a concentration of 2.5 μl per 1×10^6^ cells.

Cells were incubated for 25 minutes at room temperature in the dark. All tubes were washed once with 1 ml staining buffer, then Fixation/Permeabilization solution (250 μl) was added and cells were incubated for 20 minutes at 4°C. Cell permeability was performed using 1xBD Perm/Wash buffer, intra-cellular antibody mixtures (50 μl/Brilliant Violet 605TM anti-T-bet (BLG 644817), Eomes-conjugated-PerCP-eFluor 710 (Dan11mag) and INF-γ-conjugated-PE/VIO-770 (Miltenyi Biotec 130-109-313), CD49a− APC-Vio770 (Miltenyi Biotec 130-101-324) and FITC anti-human CD49b (BLG 359305) were added and incubated for 30 minutes at room temperature in the dark, cells were then washed twice with 1xBD Perm/Wash buffer (BD Cytofix/CytopermTM Fixation/Permeabilization Kit).

All cell samples were detected by FACS Calibur Flow Cytometer (Benton Dickinson) using Cell Quest software, and the acquired data were further analyzed using FlowJo 5.7.2 (Tree Star).

### Cytokine analyses in culture medium by ELISA

Production of IFN-γ by ILC1-like cells from healthy volunteers was analyzed using ELISA. ILC2, ILC3 and PBMCs/PHA were analyzed as negative controls. CD8+/NKG2D+ cells were analyzed as a positive control.

The concentration of IFN-γ was determined in the supernatant of 6×10^6^ cells from each donor (6 healthy donors) using the Human IFN-γ ELISA deluxe set (BioLegend) according to the manufacturer’s protocol.

### Analysis of HF cytotoxicity, catagen induction, and immune privilege collapse

As an indication of HF cytotoxicity, LDH release into the supernatants was quantified by colorimetric assay using the Cytotoxicity Detection kit Plus (Roche), which measures the conversion of tetrazolium salt in formazan, a water-soluble dye with a broad absorption maximum at approximately 500 nm (Uchida et al., 2021; Lu et al., 2007; Poeggeler et al., 2010). Medium with/without HFs was cultured with PBMCs/PHA, CD8+/NKG2D+, ILC1-like, ILC2 or ILC3 cells for three days. Formazan absorbance was measured for each condition that correlates with cell cytotoxicity. Anagen and catagen HFs were visualized and differentiated under Nikon Diaphot inverted binocular and thereafter qualitative morphological and quantitative morphometric assessments were analyzed as previously described (Kloepper et al., 2010). IHC staining was performed to test all hallmarks of AA in order to probe whether co-culture with ILC1-like cells induced abnormal HLA-DR, HLA-ABC, CD1d, ß2-microglobulin, and MICA protein expression in the proximal HF epithelium and/or downregulated the key guardians of HF immune privilege, TGF-β1 and α-MSH (Bertolini et al., 2020; Ito et al., 2004), using the qIHM method described above.

In order to check whether ILC1-like cells affect HFs via IFN-γ overproduction or via activation of the NKG2D-NKG2DL axis following excessive MICA expression by stressed HFs, neutralizing anti-IFN-γ (10 µg/ml, R&D Systems, MAB285) or function-blocking NKG2D (5 µg/ml, R&D Systems, MAB139-100) antibodies, were added to the HFs co-cultured with ILC1-like cells (defined as CD49a+ CD49b- (Verma et al., 2020), lin-/CD127+/CD117-/CRTH2-, and T-bet^lo^/ Eomes^hi^ (Bennstein et al., 2020; Krabbendam et al., 2021).

### Humanized AA mouse model

For the humanized AA mouse model (Gilhar et al., 2013a; Ghraieb et al., 2018; Gilhar et al., 2016), full-thickness biopsies were taken from healthy donors undergoing plastic surgery on the scalp. Biopsies from each donor were dissected horizontally to generate pieces with a diameter of 3 mm. Three 3 mm pieces were grafted orthotopically into the subcutaneous layer of each SCID/beige mice as previously described (Gilhar et al., 2013a; Ghraieb et al., 2018; Gilhar et al., 2013b). Seven days after surgery, mice were treated with Minoxi-5 (hair regrowth treatment for men containing 5% Minoxidil active ingredient) by spreading it on the grafts twice a day until we received optimal expedited hair growth (period of two months).

In the current study, 18 female SCID/beige mice (C.B-17/IcrHsd-scid-bg) (Harlan Laboratories Ltd., Jerusalem, Israel) were used at 2-3 months of age and were housed in the pathogen-free animal facility of the Rappaport Faculty of Medicine, Technion – Israel Institute of Technology. Animal care and research protocols were in accordance with institutional guidelines and were approved by the Institutional Committee on Animal Use (17-08-115-IL).

### Culture of Peripheral blood mononuclear cells

PBMCs were isolated from healthy donors without any history of AA or other autoimmune diseases by centrifugation on Ficoll/Hypaque (Pharmacia, Amersham Pharmacia Biotech, Uppsala, Sweden) (Ghraieb et al., 2018). The PBMCs were then cultured for 14 days with 100 U IL-2 per ml (Pepro Tech Inc, Rocky Hill, NJ) in medium composed of RPMI 1640, 10% human AB serum (Sigma, St. Louis, MO), 1% glutamine, 1% antibiotics (media components; Biological Industries, Kibbutz Beit Haemeck, Israel). Medium was changed as needed. The cultured cells defined as enriched CD8/NKG2D according to our previous publication (Ghraieb et al., 2018), were injected intradermally into human explants on beige-SCID mice.

### Study design

Two sets of experiments were performed: In the first set, the mice were divided randomly into three groups on day 89 after scalp skin transplantation and treated as described in **table supplement 2**.

The second set of experiments was performed to eliminate the confounding influence resident human T-cells present in the human scalp skin xenotransplants. To this end, anti-CD3/OKT3 antibodies (**table supplement 2**) were injected into xenotransplants treated with either autologous ILC1-like cells or autologous enriched CD8+/NKG2D+ cells. For both sets of experiments, the mice were sacrificed and skin biopsies were taken for analysis on day 45 after immunocyte injection.

### Statistical analysis

Data are presented as the mean ± standard error of mean (SEM) or fold change of mean ± SEM; p values of <0.05 were regarded as significant.

Gaussian distribution of the data was analyzed using Shapiro-Wilk test. Significant differences were analyzed using either unpaired Student′s t-test (comparison between one set of data) or One Way ANOVA (comparison between multiple sets of data) for parametric data, or Mann–Whitney test (comparison between one set of data and sham or vehicle) for nonparametric data or Kruskal–Wallis test, and Dunn’s test (comparison between multiple sets of data). The n (e.g., number of donors, tissue sections or microscopic fields) used for each individual data reported here is listed in corresponding figure legend.

## Materials and data availability statement

All data generated or analyzed during this study are included in the manuscript and supporting file; Source Data files have been provided for Figures 1–8 and figures supplement 2,3,4. Materials for all figures are provided in the Materials and Methods section.

## Acknowledgements

The study was supported in part by the Technion Research & Development Foundation (to A.G.), a Frost Endowed Scholarship from the University of Miami Dermatology Department (to R.P.), and a basic research grant from Monasterium Laboratory GmbH, Münster (to A.G. & C.R.).

## Author Contributions

A.G conceived and supervised the project. R.L.B. and A.K. performed most of the experiments. A.G. and R.P. designed the experiments, interpreted the data, and wrote the manuscript. M.B. provided the data for supplementary table 1 and edited the text and figures. Y.U. provided human skin samples. All authors edited the final manuscript.

## Online supplemental material

**Fig. S1** shows the single channels immunofluorescence microscopy analyses of various markers in AA scalp skin and in AA-induced xenotransplants.

**Fig. S2** shows the prevention of AA in scalp skin xenotransplants treated with enriched CD8/NKG2D but not with ILC1-like cells following treatment with anti-CD3 antibodies.

**Fig. S3** shows dermal infiltrates of the various treated xenotransplants.

**Fig. S4** shows the development of AA in normal human scalp skin xenotransplants treated with ILC1-like cells, enriched CD8/NKG2D and PBMCs/PHA.

**Fig. S5** shows a scheme demonstrating the isolation of ILC1-like, ILC2s and ILC3s cells from PBMCs of healthy human volunteers.

**Table. S1** shows that microdissected, organ-cultured HFs are “stressed” on day 1, but become equilibrated on day 3.

**Table. S2** shows the route of injections, volume and the number of immune cells injected into the normal healthy human xenotransplants.

## Supplementary figure legends

**Figure S1.**
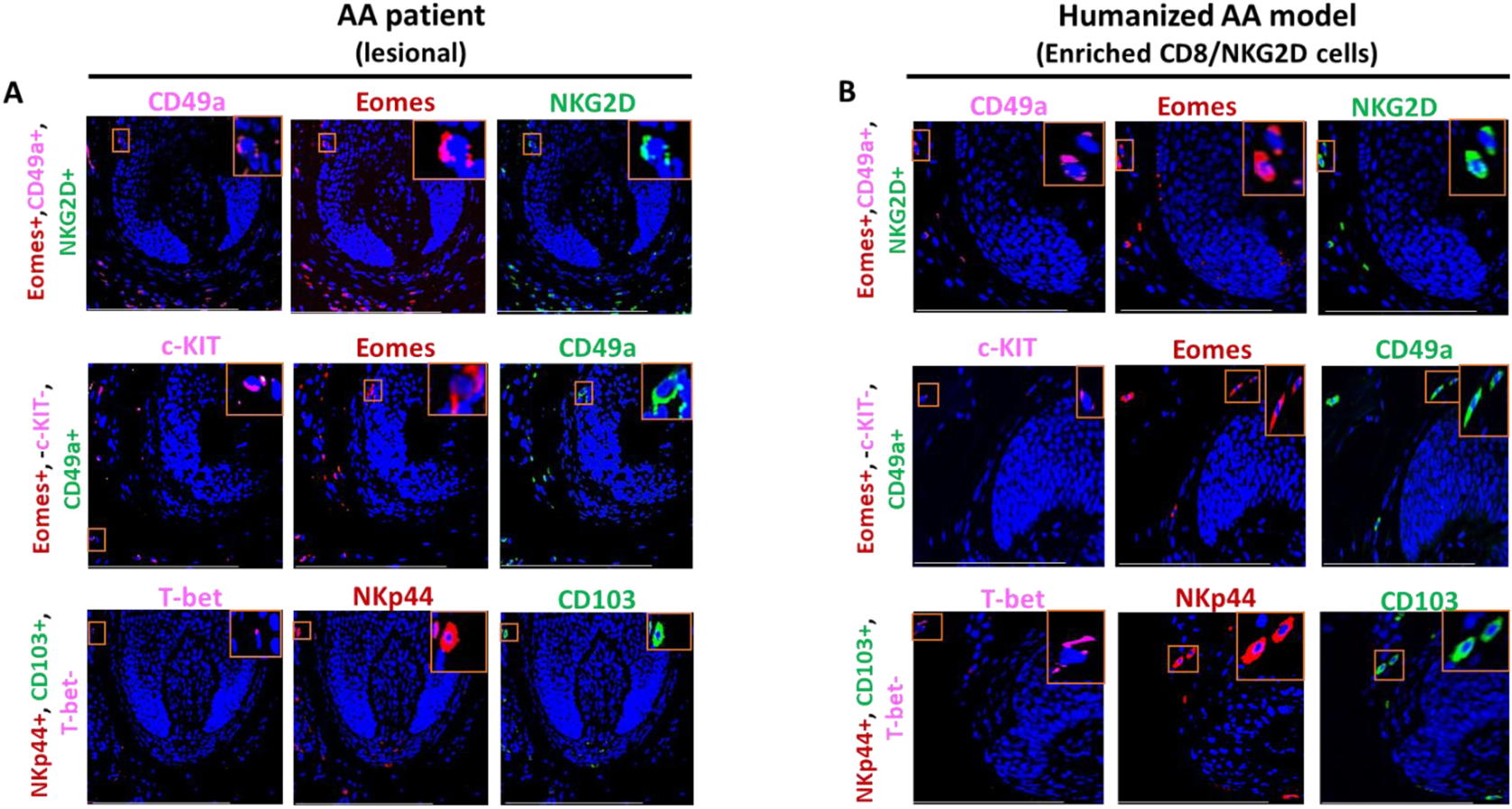
Single channels immunofluorescence microscopy analyses of various markers in AA scalp skin and in AA-induced xenotransplants. (**A**) Single channels of EOMES, T-bet, CD49a+, NKG2D+, c-kit, CD103 and NKp44 expressing cells in AA scalp patient and (**B**) in the humanized AA model. N=4 biopsies from AA patients and 5 biopsies from the humanized AA mouse model. Scale bars, 50 µm.

**Figure S2.**
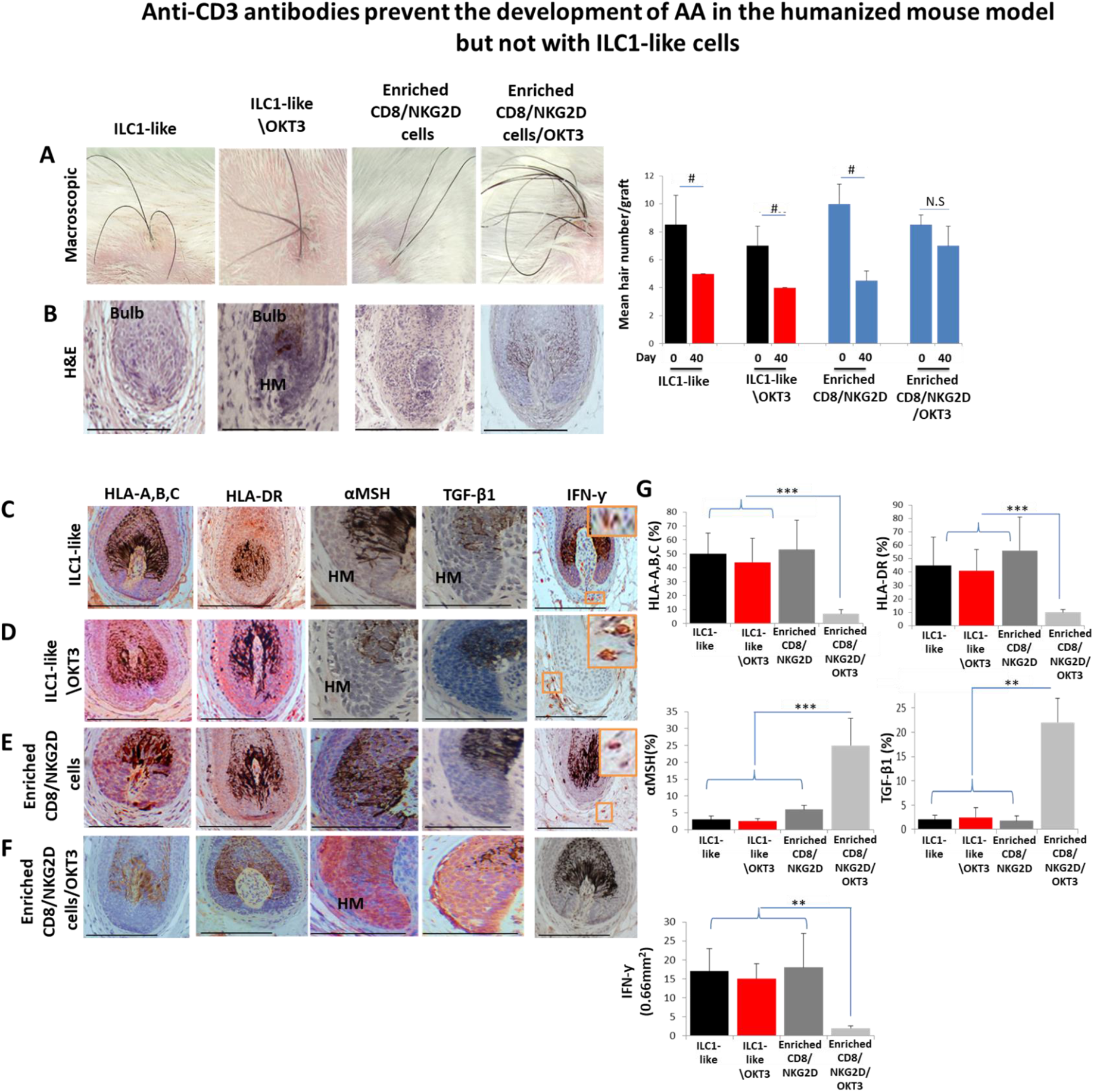
Anti-CD3 antibodies prevent the development of AA in scalp skin xenotransplants treated with enriched CD8/NKG2D but not with ILC1-like cells. (**A**) Significant hair loss was observed following the injection of ILC1-like cells, ILC1-like cells/OKT3 or enriched CD8/NKG2D cells into normal scalp skin on SCID/beige mice, but not in those treated with enriched CD8+/NKG2D+/anti-CD3 (OKT3), N=5 xenotransplants/group from 2 independent donors. Following Shapiro-Wilk test, Mann-Whitney *U* test: ^#^*p* < 0.05, ^##^*p* < 0.01. HF dystrophy combined with perifollicular lymphocytic infiltrate around anagen HFs (H&E staining) in xenotransplants treated with (**B**) ILC1-like cells, ILC1-like cells/OKT3 or enriched CD8/NKG2D. Normal HF in anagen in xenotransplant treated with enriched CD8/NKG2D/OKT3. Induction of HLA-A,B,C and of HLA-DR, downregulation of α-MSH and TGF-β1 by the follicular epithelium and increased dermal IFN-γ cells (IHC staining) in xenotransplants treated with (**C**)ILC1-like cells and similarly in (**D**) ILC1-like cells/OKT3 and (**E**) enriched CD8/NKG2D treated groups. Reduced expression of HLA-A,B,C, HLA-DR and upregulation of α-MSH and TGF-β1 combined with decreased number of dermal IFN-γ cells were observed in xenotransplants treated with (**F**) enriched CD8/NKG2D/OKT3 cells. (**G**) Quantitative data, N=5-6 xenotransplants/group from 2 independent donors. 3-4 areas were evaluated per section, and 3 sections per xenotransplant. Following Shapiro-Wilk test, Student’s *t*-test: \**p* < 0.05, \*\**p* < 0.01, \*\*\**p* < 0.001. Scale bar, 50 µm. **HM** - hair matrix.

**Figure S3.**
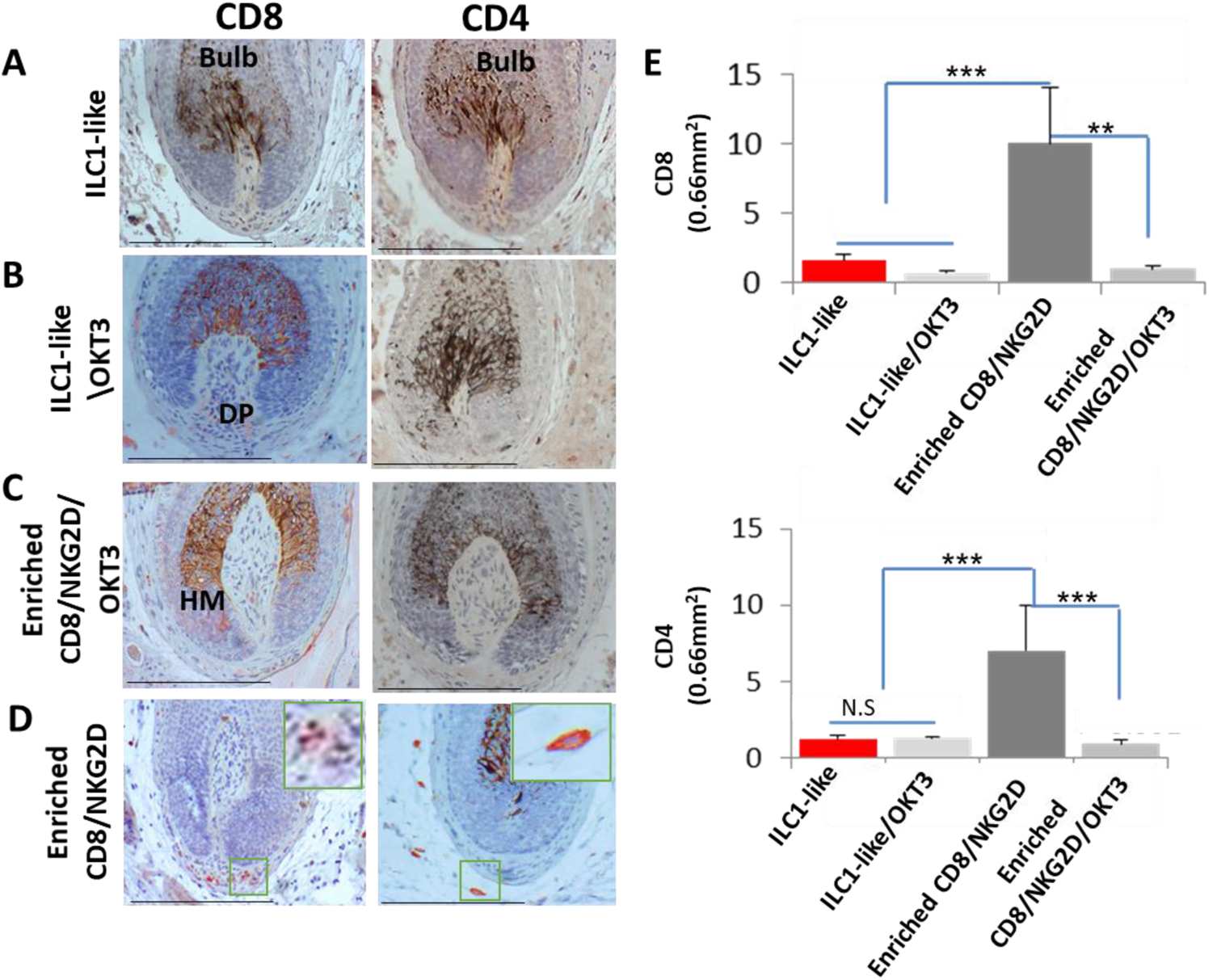
Dermal infiltrates of the various treated xenotransplants. Several CD8 and CD4 in xenotransplants treated with (**A**) ILC1s-like and similarly in xenotransplants treated with (**B**) ILC1-like cells/OKT3 and (**C**) enriched CD8/NKG2D/OKT3 cells versus increased CD8 and CD4 in xenotransplants treated with (**D**) enriched CD8/NKG2D cells. (**E**) Quantitative data, N=5-6 xenotransplants/group from 2 independent donors, 3-4 areas were evaluated per section, and 3 sections per xenotransplant. Following Shapiro-Wilk test, Student’s *t*-test: \**p* < 0.05, \*\**p* < 0.01, \*\*\**p* < 0.001. Scale bar, 50 µm. **DP** - dermal papilla, **HM** - hair matrix.

**Figure S4.**
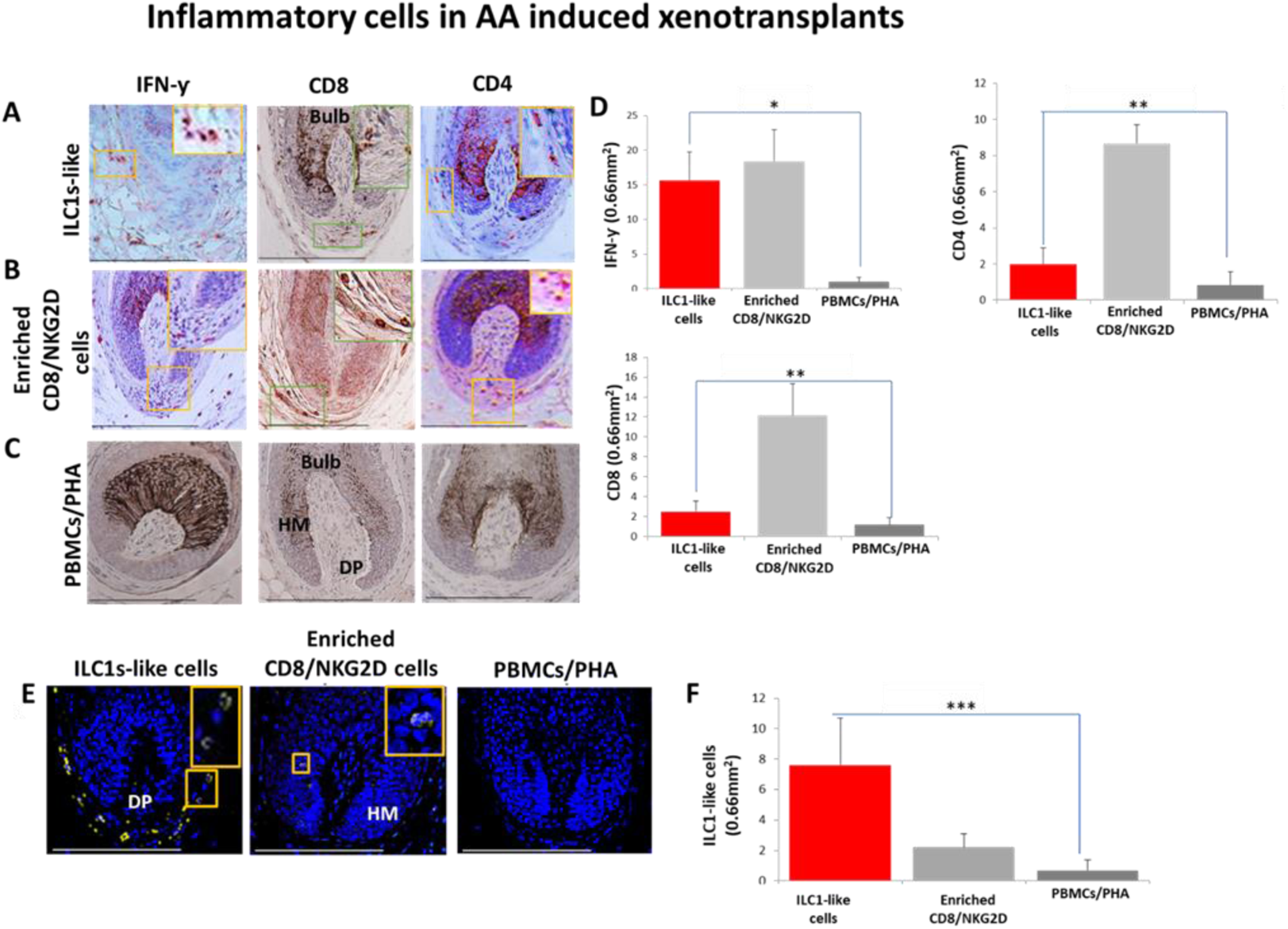
Development of AA in normal human scalp skin xenotransplants treated with (A) ILC1-like cells. Infiltrates of IFN-γ +,CD8 and CD4 cells in group treated with (**B**) enriched CD8/NKG2D cells. Decreased number of IFN-γ +, CD8 and CD4 cells in the (**C**) PBMCs/PHA treated one. (**D**) Quantitative data (**E**) presence of ILC1-like cells in transplants injected with ILC1-like cells compared to the virtual absence of these cells in enriched CD8/NKG2D and PBMCs/PHA treated mice. (**F**) Quantitative data N=5-9 xenotransplats/group from 3 donors. 4-5 areas were evaluated per section, and 3 sections per xenotransplant. Following Shapiro-Wilk test, Student’s *t*-test: \**p* < 0.05, \*\**p* < 0.01, \*\*\**p* < 0.001. Scale bar, 50 µm. **DP** - dermal papilla, **HM** - hair matrix.

**Figure S5.**
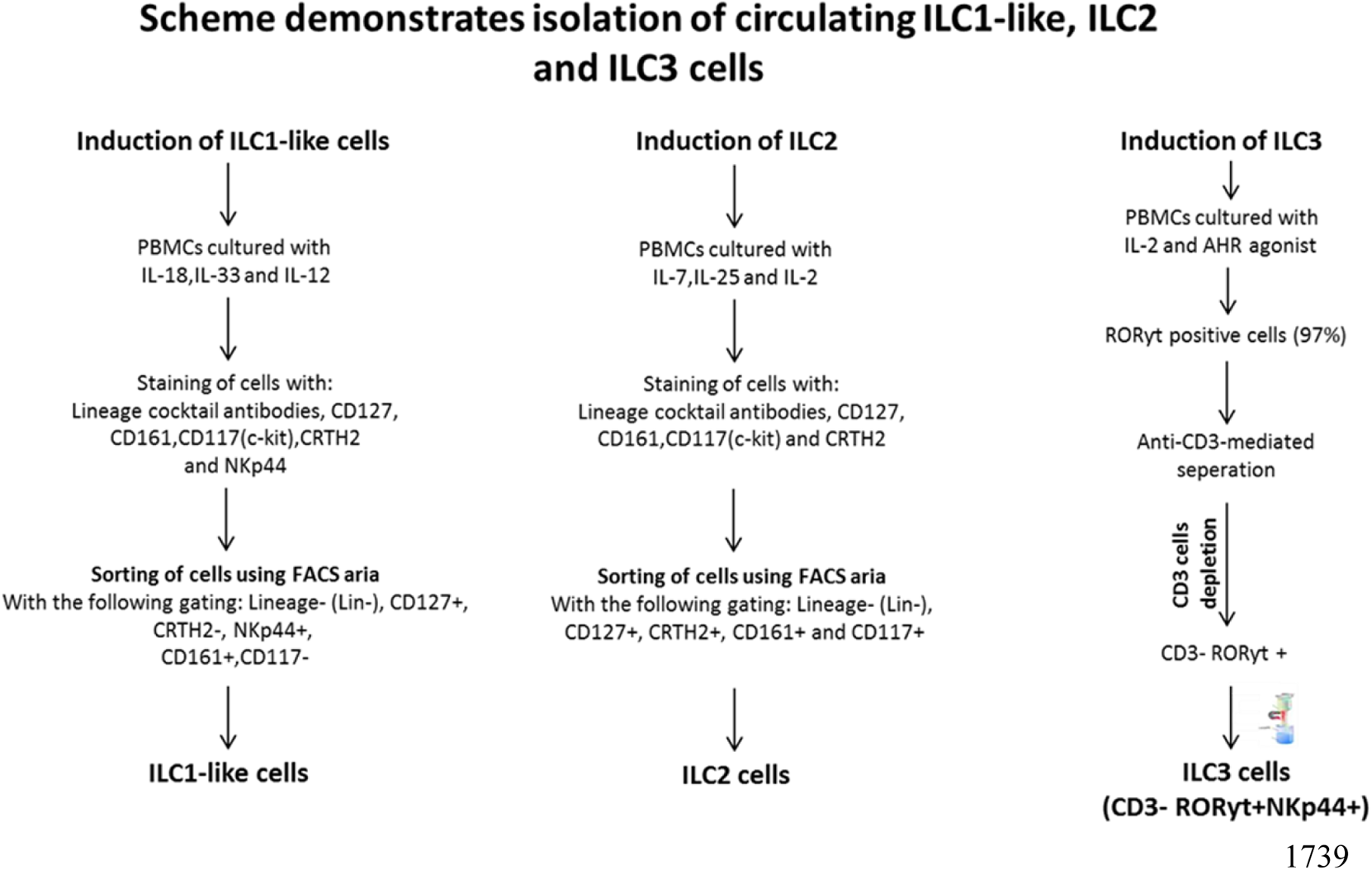
Scheme demonstrating the isolation of ILC1-like, ILC2s and ILC3s cells from PBMCs of healthy human volunteers.

**Table S1:** Microdissected, organ-cultured HFs are “stressed” on day 1, but become equilibrated on day 3, as assessed by the listed objective read-out parameters that indicate a temporarily weakened HF immune privilege and HF damage as well as MICA overexpression on day 1. Instead, the expression of HF immune privilege guardians (αMSH, TGFß2) (Bertolini et al., 2020) is preserved. This makes freshly microdissected healthy human scalp HFs one day after initiation of HF organ culture optimally suited as “stressed” human (mini-) organs that strongly express the NKG2D-activating “danger” signal”, MICA, which also is overexpressed by human AA HFs (Li et al., 2016) (these data are repeated from Uchida et al., 2021 to illustrate the HF distress/partial IP collapse of microdissected human scalp HFs one day 1 after initiation of organ culture).

| Parameter analyzed<br>Hair follicle | LDH activity | NKG2D ligand MICA | CXCL12 | IP collapse |  | IP guardians |  |
| --- | --- | --- | --- | --- | --- | --- | --- |
|  |  |  |  | MHC class I | β2-microglobulin | α-MSH | TGFβ2 |
| **Stressed**<br>(1 day in culture) | Detected | Highly detected | Detected | Detected | Detected | Detected | Highly detected |
| **Non-Stressed**<br>(3 days in culture) | Not detected | Not detected | Not detected | Not detected | Not detected | Detected | Highly detected |

\* Taken from Uchida et al., 2021

**Table S2.** Route of injections, volume and the number of immune cells injected into the normal healthy human xenotransplants.

|  | Treated group | Number of Injected Cells | Route | Injection Volume | Number of Mice |
| --- | --- | --- | --- | --- | --- |
| Experiment 1 | Enriched CD8/NKG2D cells | 4X10<sup>5</sup> cells | i.d | 100 µl | 3 |
|  | PHA activated PBMCs | 4X10<sup>5</sup> cells | i.d | 100 µl | 3 |
|  | ILC1-like cells | 4X10<sup>5</sup> cells | i.d | 100 µl | 4 |
| Experiment 2 | ILC1-like cells | 4X10<sup>5</sup> cells | i.d | 100 µl | 2 |
|  | ILC1-like cells+ anti-CD3 Abs (OKT3, Miltenyi Biotech) | 4X10<sup>5</sup> cells | i.d | 100 µl | 2 |
|  | Enriched CD8/NKG2D cells | 4X10<sup>5</sup> cells | i.d | 100 µl | 2 |
|  | Enriched CD8/NKG2D cells + anti-CD3 Abs (OKT3, Miltenyi Biotech) | 4X10<sup>5</sup> cells | i.d | 100 µl | 2 |

## References

1. Artis D, Spits H. 2015. The biology of innate lymphoid cells. Nature 15;517:293–301. doi: 10.1038/nature14189.

2. Babic M, Romagnani C. 2018. The Role of Natural Killer Group 2, Member D in Chronic Inflammation and Autoimmunity. Front Immunol 30;9:1219. doi: 10.3389/fimmu.2018.01219.

3. Bennstein SB, Weinhold S, Manser AR, Scherenschlich N, Noll A, Raba K, Kögler G, Walter L, Uhrberg M. 2020. Umbilical cord blood-derived ILC1-like cells constitute a novel precursor for mature KIR+NKG2A-NK cells. Elife 9:e55232. doi: 10.7554/eLife.55232.

4. Bertolini M, McElwee K, Gilhar A, Bulfone-Paus S, Paus R. 2020. Hair follicle immune privilege and its collapse in alopecia areata. Exp Dermatol 29:703–725. doi: 10.1111/exd.14155.

5. Bertolini M, Zilio F, Rossi A, Kleditzsch P, Emelianov VE, Gilhar A, Keren A, Meyer KC, Wang E, Funk W, McElwee K, Paus R. 2014. Abnormal interactions between perifollicular mast cells and CD8+ T-cells may contribute to the pathogenesis of alopecia areata. PLoS One 9:e94260. doi: 10.1371/journal.pone.0094260.

6. Bertolini M, Pretzlaff M, Sulk M, Bähr M, Gherardini J, Uchida Y, Reibelt M, Kinori M, Rossi A, Bíró T, Paus R. 2016. Vasoactive intestinal peptide, whose receptor-mediated signalling may be defective in alopecia areata, provides protection from hair follicle immune privilege collapse. Br J Dermatol 175:531–41. doi: 10.1111/bjd.14645.

7. Bernink JH, Mjösberg J, Spits H. 2017. Human ILC1: To Be or Not to Be. Immunity 46:756–757. doi: 10.1016/j.immuni.2017.05.001.

8. Bernink JH, Peters CP, Munneke M, te Velde AA, Meijer SL, Weijer K, Hreggvidsdottir HS, Heinsbroek SE, Legrand N, Buskens CJ, Bemelman WA, Mjösberg JM, Spits H. 2013. Human type 1 innate lymphoid cells accumulate in inflamed mucosal tissues. Nat Immunol 14:221–9. doi: 10.1038/ni.2534.

9. Bodó E, Tobin DJ, Kamenisch Y, Bíró T, Berneburg M, Funk W, Paus R. 2007.Dissecting the impact of chemotherapy on the human hair follicle: a pragmatic in vitro assay for studying the pathogenesis and potential management of hair follicle dystrophy. Am J Pathol 171:1153–67. doi: 10.2353/ajpath.2007.061164.

10. Christoph T, Müller-Röver S, Audring H, Tobin DJ, Hermes B, Cotsarelis G, Rückert R, Paus R. 2000. The human hair follicle immune system: cellular composition and immune privilege. Br J Dermatol 142:862–73. doi: 10.1046/j.1365-2133.2000.03464.x. PMID: 10809841.

11. Clottu AS, Humbel M, Fluder N, Karampetsou MP, Comte D. 2022. Innate Lymphoid Cells in Autoimmune Diseases. Front Immunol 12:789788. doi: 10.3389/fimmu.2021.789788. PMID: 35069567; PMCID: PMC8777080.

12. Colonna M. 2018. Innate Lymphoid Cells: Diversity, Plasticity, and Unique Functions in Immunity. Immunity 48:1104–1117. doi: 10.1016/j.immuni.2018.05.013. PMID: 29924976; PMCID: PMC6344351.

13. Collins A, Rothman N, Liu K, Reiner SL. 2017. Eomesodermin and T-bet mark developmentally distinct human natural killer cells. JCI Insight 2:e90063. doi: 10.1172/jci.insight.90063. PMID: 28289707; PMCID: PMC5333970.

14. Conlon TM, Knolle PA, Yildirim AÖ. 2021. Local tissue development of type 1 innate lymphoid cells: guided by interferon-gamma. Signal Transduct Target Ther 6:287. doi: 10.1038/s41392-021-00705-1. PMID: 34326313; PMCID: PMC8319591.

15. Connell SJ, Jabbari A. 2022. The current state of knowledge of the immune ecosystem in alopecia areata. Autoimmun Rev 21:103061. doi: 10.1016/j.autrev.2022.103061. Epub 2022 Feb 10. PMID: 35151885; PMCID: PMC9018517.

16. Creyns B, Jacobs I, Verstockt B, Cremer J, Ballet V, Vandecasteele R, Vanuytsel T, Ferrante M, Vermeire S, Van Assche G, Ceuppens JL, Breynaert C. 2020. Biological Therapy in Inflammatory Bowel Disease Patients Partly Restores Intestinal Innate Lymphoid Cell Subtype Equilibrium. Front Immuno 11:1847. doi: 10.3389/fimmu.2020.01847. PMID: 32983101; PMCID: PMC7481382.

17. Cruz-Zárate D, Cabrera-Rivera GL, Ruiz-Sánchez BP, Serafín-López J, Chacón-Salinas R, López-Macías C, et al. 2018.Innate Lymphoid Cells Have Decreased HLA-DR Expression but Retain Their Responsiveness to TLR Ligands during Sepsis. J Immunol 201:3401–3410. doi: 10.4049/jimmunol.1800735. Epub 2018 Oct 29. PMID: 30373848.

18. Dadi S, Chhangawala S, Whitlock BM, Franklin RA, Luo CT, Oh SA, Toure A, Pritykin Y, Huse M, Leslie CS, Li MO. 2016. Cancer Immunosurveillance by Tissue-Resident Innate Lymphoid Cells and Innate-like T Cells. Cell 164:365–77. doi: 10.1016/j.cell.2016.01.002. Epub 2016 Jan 21. PMID: 26806130; PMCID: PMC4733424.

19. Daussy C, Faure F, Mayol K, Viel S, Gasteiger G, Charrier E, Bienvenu J, Henry T, Debien E, Hasan UA, Marvel J, Yoh K, Takahashi S, Prinz I, de Bernard S, Buffat L, Walzer T. 2014. T-bet and Eomes instruct the development of two distinct natural killer cell lineages in the liver and in the bone marrow. J Exp Med 211:563–77. doi: 10.1084/jem.20131560. Epub 2014 Feb 10.

20. de Jong A, Jabbari A, Dai Z, Xing L, Lee D, Li MM, Duvic M, Hordinsky M, Norris DA, Price V, Mackay-Wiggan J, Clynes R, Christiano AM. 2018. High-throughput T cell receptor sequencing identifies clonally expanded CD8+ T cell populations in alopecia areata. JCI Insight 3:e121949. doi: 10.1172/jci.insight.121949. PMID: 30282836; PMCID: PMC6237451.

21. Ebbo M, Crinier A, Vély F, Vivier E. 2017. Innate lymphoid cells: major players in inflammatory diseases. Nat Rev Immunol 17:665–678. doi: 10.1038/nri.2017.86. Epub 2017 Aug 14. PMID: 28804130.

22. Edelkamp, J., et al. 2021. Selective Inhibition of Tyrosine Kinase 2 (TYK2) Protects Hair Follicles from Immune Privilege Collapse Induced by Interleukin (IL)-12 Stimulation. EADV’s 30th Anniversary Congress, Abstract.

23. Fan J, Shi J, Zhang Y, Liu J, An C, Zhu H, Wu P, Hu W, Qin R, Yao D, Shou X, Xu Y, Tong Z, Wen X, Xu J, Zhang J, Fang W, Lou J, Yin W, Chen W. 2022. NKG2D discriminates diverse ligands through selectively mechano-regulated ligand conformational changes. EMBO J 41:e107739. doi: 10.15252/embj.2021107739. Epub 2021 Dec 16. PMID: 34913508; PMCID: PMC8762575.

24. Fang W, Zhang Y, Chen Z. 2020. Innate lymphoid cells in inflammatory arthritis. Arthritis Res Ther 22:25. doi: 10.1186/s13075-020-2115-4. PMID: 32051038; PMCID: PMC7017550.

25. Flommersfeld S, Böttcher JP, Ersching J, Flossdorf M, Meiser P, Pachmayr LO, Leube J, Hensel I, Jarosch S, Zhang Q, Chaudhry MZ, Andrae I, Schiemann M, Busch DH, Cicin-Sain L, Sun JC, Gasteiger G, Victora GD, Höfer T, Buchholz VR, Grassmann S. 2021.Fate mapping of single NK cells identifies a type 1 innate lymphoid-like lineage that bridges innate and adaptive recognition of viral infection. Immunity 54:2288–2304.e7. doi: 10.1016/j.immuni.2021.08.002.

26. Frazao A, Rethacker L, Messaoudene M, Avril MF, Toubert A, Dulphy N, Caignard A. 2019. NKG2D/NKG2-Ligand Pathway Offers New Opportunities in Cancer Treatment. Front Immunol 10:661. doi: 10.3389/fimmu.2019.00661. PMID: 30984204; PMCID: PMC6449444.

27. Fuchs A, Vermi W, Lee JS, Lonardi S, Gilfillan S, Newberry RD, Cella M, Colonna M. 2013. Intraepithelial type 1 innate lymphoid cells are a unique subset of IL-12- and IL-15-responsive IFN-γ-producing cells. Immunity 38:769–81. doi: 10.1016/j.immuni.2013.02.010. Epub 2013 Feb 28. PMID: 23453631; PMCID: PMC3634355.

28. Gao Y, Souza-Fonseca-Guimaraes F, Bald T, Ng SS, Young A, Ngiow SF, Rautela J, Straube J, Waddell N, Blake SJ, Yan J, Bartholin L, et al. 2017. Tumor immunoevasion by the conversion of effector NK cells into type 1 innate lymphoid cells. Nat Immunol 18:1004–1015. doi: 10.1038/ni.3800. Epub 2017 Jul 31. PMID: 28759001.

29. Ghraieb A, Keren A, Ginzburg A, Ullmann Y, Schrum AG, Paus R, Gilhar A. 2018, iNKT cells ameliorate human autoimmunity: Lessons from alopecia areata. J Autoimmun 91:61–72. doi: 10.1016/j.jaut.2018.04.001. Epub 2018 Apr 18. PMID: 29680372.

30. Gilhar A, Etzioni A, Paus R. 2012. Alopecia areata. N Engl J Med 366:1515–25. doi: 10.1056/NEJMra1103442. PMID: 22512484.

31. Gilhar A, Ullmann Y, Berkutzki T, Assy B, Kalish RS. 1998. Autoimmune hair loss (alopecia areata) transferred by T lymphocytes to human scalp explants on SCID mice. J Clin Invest 101:62–7. doi: 10.1172/JCI551. PMID: 9421466; PMCID: PMC508540.

32. Gilhar A, Keren A, Shemer A, d’Ovidio R, Ullmann Y, Paus R. 2013a. Autoimmune disease induction in a healthy human organ: a humanized mouse model of alopecia areata. J Invest Dermatol 133:844–847. doi: 10.1038/jid.2012.365. Epub 2012 Oct 25. PMID: 23096715.

33. Gilhar A, Keren A, Shemer A, Ullmann Y, Paus R. 2013b. Blocking potassium channels (Kv1.3): a new treatment option for alopecia areata? J Invest Dermatol 133:2088–91. doi: 10.1038/jid.2013.141. Epub 2013 Mar 20. PMID: 23636064.

34. Gilhar A, Schrum AG, Etzioni A, Waldmann H, Paus R. 2016. Alopecia areata: Animal models illuminate autoimmune pathogenesis and novel immunotherapeutic strategies. Autoimmun Rev 15:726–35. doi: 10.1016/j.autrev.2016.03.008. Epub 2016 Mar 10. PMID: 26971464; PMCID: PMC5365233.

35. Gilhar A, Laufer-Britva R, Keren A, Paus R. 2019a. Frontiers in alopecia areata pathobiology research. J Allergy Clin Immunol 144:1478–1489. doi: 10.1016/j.jaci.2019.08.035.

36. Gilhar A, Keren A, Paus R. 2019b. JAK inhibitors and alopecia areata. Lancet 393:318–319. doi: 10.1016/S0140-6736(18)32987-8.

37. Guo C, Zhou M, Zhao S, Huang Y, Wang S, Fu R, Li M, Zhang T, Gaskin F, Yang N, Fu SM. 2019. Innate lymphoid cell disturbance with increase in ILC1 in systemic lupus erythematosus. Clin Immunol 202:49–58. doi: 10.1016/j.clim.2019.03.008. Epub 2019 Mar 26. PMID: 30926441; PMCID: PMC8191378.

38. Hawke LG, Mitchell BZ, Ormiston ML. 2020a. TGF-β and IL-15 Synergize through MAPK Pathways to Drive the Conversion of Human NK Cells to an Innate Lymphoid Cell 1-like Phenotype. J Immunol 204:3171–3181. doi: 10.4049/jimmunol.1900866. Epub 2020 Apr 24. PMID: 32332109.

39. Hawke LG, Whitford MKM, Ormiston ML. 2020b. The Production of Pro-angiogenic VEGF-A Isoforms by Hypoxic Human NK Cells Is Independent of Their TGF-β-Mediated Conversion to an ILC1-Like Phenotype. Front Immunol 11:1903. doi: 10.3389/fimmu.2020.01903. PMID: 32983113; PMCID: PMC7477355.

40. Harris JE. 2013a. Vitiligo and alopecia areata: apples and oranges? Exp Dermatol 22:785–9. doi: 10.1111/exd.12264. PMID: 24131336; PMCID: PMC3867815.

41. Harries MJ, Meyer K, Chaudhry I, E Kloepper J, Poblet E, Griffiths CE, Paus R. 2013b. Lichen planopilaris is characterized by immune privilege collapse of the hair follicle’s epithelial stem cell niche. J Pathol. 2013 231:236–47. doi: 10.1002/path.4233. PMID: 23788005.

42. Hardman-Smart JA, Purba TS, Panicker S, Farjo B, Farjo N, Harries MJ, Paus R. 2020. Does mitochondrial dysfunction of hair follicle epithelial stem cells play a role in the pathobiology of lichen planopilaris? Br J Dermatol. 183:964–966. doi: 10.1111/bjd.19259.

43. Hendrix S, Handjiski B, Peters EM, Paus R. 2005. A guide to assessing damage response pathways of the hair follicle: lessons from cyclophosphamide-induced alopecia in mice. J Invest Dermatol 125:42–51. doi: 10.1111/j.0022-202X.2005.23787.x. PMID: 15982301.

44. Ikeda T. 1965. A new classification of alopecia areata. Dermatologica 131:421–45. doi: 10.1159/000254503. PMID: 5864736.

45. Ito T, Ito N, Bettermann A, Tokura Y, Takigawa M, Paus R. 2004. Collapse and restoration of MHC class-I-dependent immune privilege: exploiting the human hair follicle as a model. Am J Pathol 164:623–34. doi: 10.1016/S0002-9440(10)63151-3. PMID: 14742267; PMCID: PMC1602279.

46. Ito T, Ito N, Saathoff M, Bettermann A, Takigawa M, Paus R. 2005a. Interferon-gamma is a potent inducer of catagen-like changes in cultured human anagen hair follicles. Br J Dermatol 152:623–31. doi: 10.1111/j.1365-2133.2005.06453.x. PMID: 15840090.

47. Ito N, Ito T, Kromminga A, Bettermann A, Takigawa M, Kees F, Straub RH, Paus R. 2005b. Human hair follicles display a functional equivalent of the hypothalamic-pituitary-adrenal axis and synthesize cortisol. FASEB J 19:1332–4. doi: 10.1096/fj.04-1968fje. Epub 2005 Jun 9. PMID: 15946990.

48. Ito T, Ito N, Saatoff M, Hashizume H, Fukamizu H, Nickoloff BJ, Takigawa M, Paus R. 2008. Maintenance of hair follicle immune privilege is linked to prevention of NK cell attack. J Invest Dermatol 128:1196–206. doi: 10.1038/sj.jid.5701183. Epub 2007 Dec 27. PMID: 18160967.

49. Ito T, Kageyama R, Nakazawa S, Honda T. 2020. Understanding the significance of cytokines and chemokines in the pathogenesis of alopecia areata. Exp Dermatol 29:726–732. doi: 10.1111/exd.14129. Epub 2020 Jul 3. PMID: 32533873.

50. Jiao Y, Wu L, Huntington ND, Zhang X. 2020. Crosstalk Between Gut Microbiota and Innate Immunity and Its Implication in Autoimmune Diseases. Front Immunol 11:282. doi: 10.3389/fimmu.2020.00282. PMID: 32153586; PMCID: PMC7047319.

51. Jiao Y, Huntington ND, Belz GT, Seillet C. 2016. Type 1 Innate Lymphoid Cell Biology: Lessons Learnt from Natural Killer Cells. Front Immunol 7:426. doi: 10.3389/fimmu.2016.00426. PMID: 27785129; PMCID: PMC5059362.

52. Keren A, Shemer A, Ullmann Y, Paus R, Gilhar A. 2014. The PDE4 inhibitor, apremilast, suppresses experimentally induced alopecia areata in human skin in vivo. J Dermatol Sci 77:74–6. doi: 10.1016/j.jdermsci.2014.11.009. Epub 2014 Dec 3. PMID: 25530115.

53. Keren A, Shemer A, Ginzburg A, Ullmann Y, Schrum AG, Paus R, Gilhar A. 2018. Innate lymphoid cells 3 induce psoriasis in xenotransplanted healthy human skin. J Allergy Clin Immunol 142:305–308.e6. doi: 10.1016/j.jaci.2018.02.015. Epub 2018 Mar 2. PMID: 29501801.

54. Kim J, Ryu S, Kim HY. 2021. Innate Lymphoid Cells in Tissue Homeostasis and Disease Pathogenesis. Mol Cells 44:301–309. doi: 10.14348/molcells.2021.0053. PMID: 33972473; PMCID: PMC8175152.

55. Kim BS. 2015. Innate lymphoid cells in the skin. J Invest Dermatol 135:673–678. doi: 10.1038/jid.2014.401. Epub 2014 Oct 23. PMID: 25339380.

56. King BA, Mesinkovska NA, Craiglow B, Kindred C, Ko J, McMichael A, Shapiro J, Goh C, Mirmirani P, Tosti A, Hordinsky M, Huang KP, Castelo-Soccio L, Bergfeld W, Paller AS, Mackay-Wiggan J, Glashofer M, et al. 2022. Development of the alopecia areata scale for clinical use: Results of an academic-industry collaborative effort. J Am Acad Dermatol 86:359–364. doi: 10.1016/j.jaad.2021.08.043. Epub 2021 Aug 30. PMID: 34474079.

57. Kinori M, Bertolini M, Funk W, Samuelov L, Meyer KC, Emelianov VU, Hasse S, Paus R. 2012. Calcitonin gene-related peptide (CGRP) may award relative protection from interferon-γ-induced collapse of human hair follicle immune privilege. Exp Dermatol 21:223–226. doi: 10.1111/j.1600-0625.2011.01432.x. PMID: 22379970.

58. Kloepper JE, Sugawara K, Al-Nuaimi Y, Gáspár E, van Beek N, Paus R. 2010. Methods in hair research: how to objectively distinguish between anagen and catagen in human hair follicle organ culture. Exp Dermatol 19:305–12. doi: 10.1111/j.1600-0625.2009.00939.x. Epub 2009 Aug 31. PMID: 19725870.

59. Korta DZ, Christiano AM, Bergfeld W, Duvic M, Ellison A, Fu J, Harris JE, Hordinsky MK, King B, Kranz D, Mackay-Wiggan J, McMichael A, Norris DA, Price V, Shapiro J, Atanaskova Mesinkovska N. 2018. Alopecia areata is a medical disease. J Am Acad Dermatol 78:832–834. doi: 10.1016/j.jaad.2017.09.011. PMID: 29548423.

60. Krabbendam L, Bernink JH, Spits H. 2021. Innate lymphoid cells: from helper to killer. Curr Opin Immunol 68:28–33. doi: 10.1016/j.coi.2020.08.007. Epub 2020 Sep 21. PMID: 32971468.

61. Krueger JG, McInnes IB, Blauvelt A. 2022. Tyrosine kinase 2 and Janus kinase‒ signal transducer and activator of transcription signaling and inhibition in plaque psoriasis. J Am Acad Dermatol 86:148–157. doi: 10.1016/j.jaad.2021.06.869. Epub 2021 Jul 2. PMID: 34224773.

62. Langan EA, Philpott MP, Kloepper JE, Paus R. 2015. Human hair follicle organ culture: theory, application and perspectives. Exp Dermatol 24:903–11. doi: 10.1111/exd.12836. PMID: 26284830.

63. Laufer Britva R, Keren A, Paus R, Gilhar A. 2020. Apremilast and tofacitinib exert differential effects in the humanized mouse model of alopecia areata. Br J Dermatol 182:227–229. doi: 10.1111/bjd.18264. Epub 2019 Sep 5. PMID: 31254391.

64. Li J, van Vliet C, Rufaut NW, Jones LN, Sinclair RD, Carbone FR. 2016. Laser Capture Microdissection Reveals Transcriptional Abnormalities in Alopecia Areata before, during, and after Active Hair Loss. J Invest Dermatol 136:715–718. doi: 10.1016/j.jid.2015.12.003. Epub 2015 Dec 10. PMID: 27015457.

65. Lousada MB, Lachnit T, Edelkamp J, Rouillé T, Ajdic D, Uchida Y, Di Nardo A, Bosch TCG, Paus R. 2021. Exploring the human hair follicle microbiome. Br J Dermatol 184:802–815. doi: 10.1111/bjd.19461. Epub 2021 Feb 18. PMID: 32762039.

66. Lu Z, Hasse S, Bodo E, Rose C, Funk W, Paus R. 2007. Towards the development of a simplified long-term organ culture method for human scalp skin and its appendages under serum-free conditions. Exp Dermatol 16:37–44. doi: 10.1111/j.1600-0625.2006.00510.x. PMID: 17181635.

67. Luo W, Tian L, Tan B, Shen Z, Xiao M, Wu S, Meng X, Wu X, Wang X. 2022. Update: Innate Lymphoid Cells in Inflammatory Bowel Disease. Dig Dis Sci 67:56–66. doi: 10.1007/s10620-021-06831-8. Epub 2021 Feb 20. PMID: 33609209.

68. McDonald BD, Jabri B, Bendelac A. 2018. Diverse developmental pathways of intestinal intraepithelial lymphocytes. Nat Rev Immunol 18:514–525. doi: 10.1038/s41577-018-0013-7. PMID: 29717233; PMCID: PMC6063796.

69. Meah N, Wall D, York K, Bhoyrul B, Bokhari L, Sigall DA, Bergfeld WF, Betz RC, Blume-Peytavi U, Callender V, Chitreddy V, Combalia A, Cotsarelis G, Craiglow B, Donovan J, Eisman S, et al. 2020. The Alopecia Areata Consensus of Experts (ACE) study: Results of an international expert opinion on treatments for alopecia areata. J Am Acad Dermatol 83:123–130. doi: 10.1016/j.jaad.2020.03.004. Epub 2020 Mar 9. PMID: 32165196.

70. Meah N, Wall D, York K, Bhoyrul B, Bokhari L, Asz-Sigall D, Bergfeld WF, Betz RC, Blume-Peytavi U, Callender V, Chitreddy V, et al. 2021.The Alopecia Areata Consensus of Experts (ACE) study part II: Results of an international expert opinion on diagnosis and laboratory evaluation for alopecia areata. J Am Acad Dermatol 84:1594–1601. doi: 10.1016/j.jaad.2020.09.028. Epub 2020 Sep 12. PMID: 32926985.

71. Messenger AG, Slater DN, Bleehen SS. 1986. Alopecia areata: alterations in the hair growth cycle and correlation with the follicular pathology. Br J Dermatol 114:337–47. doi: 10.1111/j.1365-2133.1986.tb02825.x. PMID: 3954954.

72. Mjösberg JM, Trifari S, Crellin NK, Peters CP, van Drunen CM, Piet B, Fokkens WJ, Cupedo T, Spits H. 2011. Human IL-25- and IL-33-responsive type 2 innate lymphoid cells are defined by expression of CRTH2 and CD161. Nat Immunol 12:1055–62. doi: 10.1038/ni.2104. PMID: 21909091.

73. Mora-Velandia LM, Castro-Escamilla O, Méndez AG, Aguilar-Flores C, Velázquez-Avila M, Tussié-Luna MI, Téllez-Sosa J, Maldonado-García C, Jurado-Santacruz F, Ferat-Osorio E, Martínez-Barnetche J, Pelayo R, Bonifaz LC. 2017. A Human Lin-CD123+ CD127low Population Endowed with ILC Features and Migratory Capabilities Contributes to Immunopathological Hallmarks of Psoriasis. Front Immunol 8:176. doi: 10.3389/fimmu.2017.00176. PMID: 28303135; PMCID: PMC5332395.

74. Nabekura T, Shibuya A. 2021a. Type 1 innate lymphoid cells: Soldiers at the front line of immunity. Biomed J 44:115–122. doi: 10.1016/j.bj.2020.10.001. Epub 2020 Nov 19. PMID: 33839081; PMCID: PMC8178574.

75. Nabekura T, Shibuya A. 2021b. ILC1: guardians of the oral mucosa against enemy viruses. Immunity 54:196–198. doi: 10.1016/j.immuni.2021.01.002. PMID: 33567258; PMCID: PMC7871888.

76. Nagasawa M, Heesters BA, Kradolfer CMA, Krabbendam L, Martinez-Gonzalez I, de Bruijn MJW, Golebski K, Hendriks RW, Stadhouders R, Spits H, Bal SM. 2019. KLRG1 and NKp46 discriminate subpopulations of human CD117+CRTH2-ILCs biased toward ILC2 or ILC3. J Exp Med 216:1762–1776. doi: 10.1084/jem.20190490. Epub 2019 Jun 14. Erratum in: J Exp Med. 2019 Aug 6;: PMID: 31201208; PMCID: PMC6683990.

77. Ohne Y, Silver JS, Thompson-Snipes L, Collet MA, Blanck JP, Cantarel BL, Copenhaver AM, Humbles AA, Liu YJ. 2016. IL-1 is a critical regulator of group 2 innate lymphoid cell function and plasticity. Nat Immunol 17:646–55. doi: 10.1038/ni.3447. Epub 2016 Apr 25. Erratum in: Nat Immunol. 2016 Jul 19;17(8):1005. PMID: 27111142.

78. Orimo K, Saito H, Matsumoto K, Morita H. 2020. Innate Lymphoid Cells in the Airways: Their Functions and Regulators. Allergy Asthma Immunol Res 12:381–398. doi: 10.4168/aair.2020.12.3.381. PMID: 32141254; PMCID: PMC7061164.

79. Park E, Patel S, Wang Q, Andhey P, Zaitsev K, Porter S, Hershey M, Bern M, Plougastel-Douglas B, Collins P, Colonna M, Murphy KM, Oltz E, Artyomov M, Sibley LD, Yokoyama WM. 2019. Toxoplasma gondii infection drives conversion of NK cells into ILC1-like cells. Elife 8:e47605. doi: 10.7554/eLife.47605. PMID: 31393266; PMCID: PMC6703900.

80. Paus R, Slominski A, Czarnetzki BM. 1993. Is alopecia areata an autoimmune-response against melanogenesis-related proteins, exposed by abnormal MHC class I expression in the anagen hair bulb? Yale J Biol Med 66:541–54. PMID: 7716973; PMCID: PMC2588848.

81. Paus R, Nickoloff BJ, Ito T. 2005. A ‘hairy’ privilege. Trends Immunol 26:32–40. doi: 10.1016/j.it.2004.09.014. PMID: 15629407.

82. Paus R, Bulfone-Paus S, Bertolini M. 2018. Hair Follicle Immune Privilege Revisited: The Key to Alopecia Areata Management. J Investig Dermatol Symp Proc 19:S12–S17. doi: 10.1016/j.jisp.2017.10.014. PMID: 29273098.

83. Paus R. 2020. The Evolving Pathogenesis of Alopecia Areata: Major Open Questions. J Investig Dermatol Symp Proc 20:S6–S10. doi: 10.1016/j.jisp.2020.04.002. PMID: 33099388.

84. Peters EM, Liotiri S, Bodó E, Hagen E, Bíró T, Arck PC, Paus R. 2007. Probing the effects of stress mediators on the human hair follicle: substance P holds central position. Am J Pathol 171:1872–86. doi: 10.2353/ajpath.2007.061206. Epub 2007 Nov 30. PMID: 18055548; PMCID: PMC2111110.

85. Peters EM, Stieglitz MG, Liezman C, Overall RW, Nakamura M, Hagen E, Klapp BF, Arck P, Paus R. 2006. p75 Neurotrophin Receptor-Mediated Signaling Promotes Human Hair Follicle Regression (Catagen). Am J Pathol 168:221–34. doi: 10.2353/ajpath.2006.050163. PMID: 16400025; PMCID: PMC1592649.

86. Petukhova L, Duvic M, Hordinsky M, Norris D, Price V, Shimomura Y, Kim H, Singh P, Lee A, Chen WV, Meyer KC, Paus R, Jahoda CA, Amos CI, Gregersen PK, Christiano AM. 2010. Genome-wide association study in alopecia areata implicates both innate and adaptive immunity. Nature 466:113–7. doi: 10.1038/nature09114. PMID: 20596022; PMCID: PMC2921172.

87. Pinto D, Calabrese FM, De Angelis M, Celano G, Giuliani G, Gobbetti M, Rinaldi F. 2020. Predictive Metagenomic Profiling, Urine Metabolomics, and Human Marker Gene Expression as an Integrated Approach to Study Alopecia Areata. Front Cell Infect Microbiol 10:146. doi: 10.3389/fcimb.2020.00146. PMID: 32411613; PMCID: PMC7201066.

88. Poeggeler B, Bodó E, Nadrowitz R, Dunst J, Paus R. 2010. A simple assay for the study of human hair follicle damage induced by ionizing irradiation. Exp Dermatol 19:e306–9. doi: 10.1111/j.1600-0625.2009.01009.x. PMID: 19925637.

89. Pratt CH, King LE Jr, Messenger AG, Christiano AM, Sundberg JP. 2017. Alopecia areata. Nat Rev Dis Primers 3:17011. doi: 10.1038/nrdp.2017.11. PMID: 28300084; PMCID: PMC5573125.

90. Purba TS, Brunken L, Hawkshaw NJ, Peake M, Hardman J, Paus R. 2016. A primer for studying cell cycle dynamics of the human hair follicle. Exp Dermatol 25:663–8. doi: 10.1111/exd.13046. Epub 2016 Jul 18. PMID: 27094702.

91. Quintino-de-Carvalho IL, Gonçalves-Pereira MH, Faria Ramos M, de Aguiar Milhim BHG, Da Costa ÚL, Santos ÉG, Nogueira ML, Da Costa Santiago H. 2022. Type 1 Innate Lymphoid Cell and Natural Killer Cells Are Sources of Interferon-γ and Other Inflammatory Cytokines Associated With Distinct Clinical Presentation in Early Dengue Infection. J Infect Dis 225:84–93. doi: 10.1093/infdis/jiab312. PMID: 34125227.

92. Resende M, Cardoso MS, Ribeiro AR, Flórido M, Borges M, Castro AG, Alves NL, Cooper AM, Appelberg R. 2017. Innate IFN-γ-Producing Cells Developing in the Absence of IL-2 Receptor Common γ-Chain. J Immunol 199:1429–1439. doi: 10.4049/jimmunol.1601701. Epub 2017 Jul 7. PMID: 28687660.

93. Rose NR, Bona C. 1993. Defining criteria for autoimmune diseases (Witebsky’s postulates revisited). Immunol Today 14:426–30. doi: 10.1016/0167-5699(93)90244-F. PMID: 8216719.

94. Salimi M, Ogg G. 2014. Innate lymphoid cells and the skin. BMC Dermatol 14:18. doi: 10.1186/1471-5945-14-18. PMID: 25427661; PMCID: PMC4289267.

95. Seillet C, Brossay L, Vivier E. 2021. Natural killers or ILC1s? That is the question. Curr Opin Immunol 68:48–53. doi: 10.1016/j.coi.2020.08.009. Epub 2020 Oct 14. PMID: 33069142; PMCID: PMC7925336.

96. Shannon JP, Vrba SM, Reynoso GV, Wynne-Jones E, Kamenyeva O, Malo CS, Cherry CR, McManus DT, Hickman HD. 2020. Group 1 innate lymphoid-cell-derived interferon-γ maintains anti-viral vigilance in the mucosal epithelium. Immunity 54:276–290.e5. doi: 10.1016/j.immuni.2020.12.004. Epub 2021 Jan 11. PMID: 33434494; PMCID: PMC7881522.

97. Silver JS, Kearley J, Copenhaver AM, Sanden C, Mori M, Yu L, Pritchard GH, Berlin AA, Hunter CA, Bowler R, Erjefalt JS, Kolbeck R, Humbles AA. 2016. Inflammatory triggers associated with exacerbations of COPD orchestrate plasticity of group 2 innate lymphoid cells in the lungs. Nat Immunol 17:626–35. doi: 10.1038/ni.3443. Epub 2016 Apr 25.

98. Simoni Y, Newell EW. 2017. Toward Meaningful Definitions of Innate-Lymphoid-Cell Subsets. Immunity 46:760–761. doi: 10.1016/j.immuni.2017.04.026. PMID: 28514678.

99. Spits H, Bernink JH, Lanier L. 2016. NK cells and type 1 innate lymphoid cells: partners in host defense. Nat Immunol 17:758–64. doi: 10.1038/ni.3482. PMID: 27328005.

100. Talayero P, Mancebo E, Calvo-Pulido J, Rodríguez-Muñoz S, Bernardo I, Laguna-Goya R, Cano-Romero FL, García-Sesma A, Loinaz C, Jiménez C, Justo I, Paz-Artal E. 2016. Innate Lymphoid Cells Groups 1 and 3 in the Epithelial Compartment of Functional Human Intestinal Allografts. Am J Transplant 16:72–82. doi: 10.1111/ajt.13435. Epub 2015 Aug 28. PMID: 26317573.

101. Tang Y, Tan SA, Iqbal A, Li J, Glover SC. 2019. STAT3 Genotypic Variant rs744166 and Increased Tyrosine Phosphorylation of STAT3 in IL-23 Responsive Innate Lymphoid Cells during Pathogenesis of Crohn’s Disease. J Immunol Res 2019:9406146. doi: 10.1155/2019/9406146. PMID: 31321245; PMCID: PMC6610725.

102. Teunissen MBM, Munneke JM, Bernink JH, Spuls PI, Res PCM, Te Velde A, Cheuk S, Brouwer MWD, Menting SP, Eidsmo L, Spits H, Hazenberg MD, Mjösberg J. 2014. Composition of innate lymphoid cell subsets in the human skin: enrichment of NCR(+) ILC3 in lesional skin and blood of psoriasis patients. J Invest Dermatol 134:2351–2360. doi: 10.1038/jid.2014.146. Epub 2014 Mar 21. PMID: 24658504.

103. Tomaszewska K, Kozłowska M, Kaszuba A, Lesiak A, Narbutt J, Zalewska-Janowska A. 2020. Increased Serum Levels of IFN-γ, IL-1β, and IL-6 in Patients with Alopecia Areata and Nonsegmental Vitiligo. Oxid Med Cell Longev 2020:5693572. doi: 10.1155/2020/5693572. PMID: 32832001; PMCID: PMC7421748.

104. Tulic MK, Cavazza E, Cheli Y, Jacquel A, Luci C, Cardot-Leccia N, Hadhiri-Bzioueche H, Abbe P, Gesson M, Sormani L, Regazzetti C, Beranger GE, Lereverend C, Pons C, Khemis A, Ballotti R, Bertolotto C, Rocchi S, Passeron T. 2019. Innate lymphocyte-induced CXCR3B-mediated melanocyte apoptosis is a potential initiator of T-cell autoreactivity in vitiligo. Nat Commun 10:2178. doi: 10.1038/s41467-019-09963-8. PMID: 31097717; PMCID: PMC6522502.

105. Uchida Y, Gherardini J, Schulte-Mecklenbeck A, Alam M, Chéret J, Rossi A, Kanekura T, Gross CC, Arakawa A, Gilhar A, Bertolini M, Paus R. 2020. Pro-inflammatory Vδ1+T-cells infiltrates are present in and around the hair bulbs of non-lesional and lesional alopecia areata hair follicles. J Dermatol Sci 100:129–138. doi: 10.1016/j.jdermsci.2020.09.001. Epub 2020 Sep 18. PMID: 33039243.

106. Uchida Y, Gherardini J, Pappelbaum K, Chéret J, Schulte-Mecklenbeck A, Gross CC, Strbo N, Gilhar A, Rossi A, Funk W, Kanekura T, Almeida L, Bertolini M, Paus R. 2021. Resident human dermal γδT-cells operate as stress-sentinels: Lessons from the hair follicle. J Autoimmun 124:102711. doi: 10.1016/j.jaut.2021.102711. Epub 2021 Aug 31. PMID: 34479087.

107. Ullrich KA, Schulze LL, Paap EM, Müller TM, Neurath MF, Zundler S. 2020. Immunology of IL-12: An update on functional activities and implications for disease. EXCLI J 19:1563–1589. doi: 10.17179/excli2020-3104. PMID: 33408595; PMCID: PMC7783470.

108. Verma R, Er JZ, Pu RW, Sheik Mohamed J, Soo RA, Muthiah HM, Tam JKC, Ding JL. 2020. Eomes Expression Defines Group 1 Innate Lymphoid Cells During Metastasis in Human and Mouse. Front Immunol 11:1190. doi: 10.3389/fimmu.2020.01190. PMID: 32625207; PMCID: PMC7311635.

109. Vienne M, Etiennot M, Escalière B, Galluso J, Spinelli L, Guia S, Fenis A, Vivier E, Kerdiles YM. 2021. Type 1 Innate Lymphoid Cells Limit the Antitumoral Immune Response. Front Immunol 12:768989. doi: 10.3389/fimmu.2021.768989. PMID: 34868026; PMCID: PMC8637113.

110. Vivier E, Artis D, Colonna M, Diefenbach A, Di Santo JP, Eberl G, Koyasu S, Locksley RM, McKenzie ANJ, Mebius RE, Powrie F, Spits H. 2018. Innate Lymphoid Cells: 10 Years On. Cell 174:1054–1066. doi: 10.1016/j.cell.2018.07.017. PMID: 30142344.

111. Vivier E. 2021. The discovery of innate lymphoid cells. Nat Rev Immunol 21:616. doi: 10.1038/s41577-021-00595-y. PMID: 34580448.

112. Wang EHC, Yu M, Breitkopf T, Akhoundsadegh N, Wang X, Shi FT, Leung G, Dutz JP, Shapiro J, McElwee KJ. 2016. Identification of Autoantigen Epitopes in Alopecia Areata. J Invest Dermatol 136:1617–1626. doi: 10.1016/j.jid.2016.04.004. Epub 2016 Apr 16. PMID: 27094591.

113. Wu C, He S, Liu J, Wang B, Lin J, Duan Y, Gao X, Li D. 2018. Type 1 innate lymphoid cell aggravation of atherosclerosis is mediated through TLR4. Scand J Immunol 87:e12661. doi: 10.1111/sji.12661. PMID: 29570822.

114. Xing L, Dai Z, Jabbari A, Cerise JE, Higgins CA, Gong W, de Jong A, Harel S, DeStefano GM, Rothman L, Singh P, Petukhova L, Mackay-Wiggan J, Christiano AM, Clynes R. 2014. Alopecia areata is driven by cytotoxic T lymphocytes and is reversed by JAK inhibition. Nat Med 20:1043–9. doi: 10.1038/nm.3645. Epub 2014 Aug 17. PMID: 25129481; PMCID: PMC4362521.

115. Yang Z, Tang T, Wei X, Yang S, Tian Z. 2015. Type 1 innate lymphoid cells contribute to the pathogenesis of chronic hepatitis B. Innate Immun 2:665–73. doi: 10.1177/1753425915586074. Epub 2015 May 14. PMID: 25977358.

116. Zhou S, Li Q, Wu H, Lu Q. 2020. The pathogenic role of innate lymphoid cells in autoimmune-related and inflammatory skin diseases. Cell Mol Immunol 17:335–346. doi: 10.1038/s41423-020-0399-6. Epub 2020 Mar 19. PMID: 32203190; PMCID: PMC7109064.

117. Zhang J, Marotel M, Fauteux-Daniel S, Mathieu AL, Viel S, Marçais A, Walzer T. 2018. T-bet and Eomes govern differentiation and function of mouse and human NK cells and ILC1. Eur J Immunol 48:738–750. doi: 10.1002/eji.201747299. Epub 2018 Feb 28. PMID: 29424438.

118. Zook EC, Kee BL. 2016. Development of innate lymphoid cells. Nat Immunol 17:775–82. doi: 10.1038/ni.3481. PMID: 27328007.

